# The initial response of females towards congeneric males matches the propensity to hybridise in *Ophthalmotilapia*

**DOI:** 10.1101/2021.08.07.455508

**Authors:** Maarten Van Steenberge, Noémie Jublier, Loïc Kéver, Sophie Gresham, Sofie Derycke, Jos Snoeks, Eric Parmentier, Pascal Poncin, Erik Verheyen

## Abstract

Cichlid radiations often harbour closely related species with overlapping niches and distribution ranges. Such species sometimes hybridise in nature, which raises the question how can they coexist. This also holds for the Tanganyika mouthbrooders *Ophthalmotilapia ventralis* and *O. nasuta*. Earlier studies found indications of asymmetrical hybridisation with females of *O. ventralis* accepting males of *O. nasuta*, but not the other way around. We hypothesised that this was due to differences in the capacity for species recognition. Given the higher propensity of *O. ventralis* females towards hybridisation, we expect a reduced ability for species recognition in *O. ventralis* females, compared to *O. nasuta* females. We staged two experiments, one focusing on 22 female *O. nasuta* and one on 21 female *O. ventralis*. These fish were placed in one half of a tank and briefly exposed to a conspecific or a heterospecific male, a conspecific female, or nothing (control). Female response was evaluated by scoring six tracking parameters and by noting the occurrence of ten discrete behaviours before and during the encounter. Females always responded to the presence of another fish by approaching it. Remarkably, for both *O. nasuta* and *O. ventralis*, we did not find a different response between encounters with conspecific males and females. However, in agreement with our hypothesis, *O. nasuta* females behaved differently towards conspecific or heterospecific males, whereas *O. ventralis* females did not. When presented with a heterospecific male, *O. nasuta* females performed a lower number of ‘ram’ behaviours. Additionally, they never displayed the ‘flee’ behaviour, a component of the species’ mating repertoire that was seen in all but one of the presentations with a conspecific male. Our findings show that differences in species recognition at first encounter predict to a large degree the outcome of the mating process, even in the absence of mating behaviour.

## Introduction

Speciation is traditionally seen as a gradual build-up of reproductive isolation between diverging populations (Mayr 1982; Coyne and Orr 2004). The classical view that this is a slow process that occurs between allopatric populations has recently been challenged by genomic findings (Marques et al. 2019) that showed how hybridisation can drive rapid speciation (Seehausen 2004; Jiggins et al. 2008). However, as unrestrained gene flow inevitably homogenizes the genomes of diverging lineages (Roux et al. 2016), the question remains what mechanisms keep incipient species separated. In scenarios of sympatric, closely related species, the ability to correctly distinguish between conspecific and heterospecific mates is probably crucial to maintain the integrity of the species (Sullivan 2009).

Mating is the end point of a complex decision-making process in which several potential mates are evaluated (Luttbeg et al. 2001). An encounter with a potential mate can be seen as the first step in this process. The outcome of this initial contact can have profound implications on fitness, either through the refusal of suitable mates, or through the acceptance of suboptimal partners. Therefore, we expect intra- and interspecific differences in individual responses when confronted with a choice of partners. Additionally, the preference for a given mate can depend on environmental, social and intrinsic parameters, explaining variation in preference both between and within species, and between and within individuals (Pfennig 2008; Sommer-Trembo et al. 2017). Assortative mate selection was traditionally seen as a sequential process in which individuals first assess whether ‘the other’ is a conspecific and then assess its quality as a mate (Mayr 1982). However, empirical data and theory suggest that assessment of species and quality are not independent steps (Sullivan 2009; Mendelson and Shaw 2012) as the specific status of an individual could be judged using the same cues as its quality. Additionally, adaptive hybridisation has been observed in several taxa (overview in Mendelson and Shaw 2012), indicating that preferred mates are not necessarily always conspecific.

Cichlids of the large East African Lakes form endemic, species-rich radiations (Salzburger 2018). Several suggested “key adaptations” of cichlids, such as their pharyngeal jaws, help to explain their evolution into numerous trophic niches (Kocher 2004). However, a large proportion of these closely related species coexist without apparent eco-morphological differences (Van Oppen et al. 1998). Since most cichlid assemblages are relatively young, several taxa may be classified as incipient species and they retain the potential to hybridise. The oldest East African lake, Lake Tanganyika, however, contains a mature cichlid radiation (Salzburger 2018) in which most species are well-delineated (Ronco et al. 2020). However, even between well-delineated biological species, boundaries can be permeable and molecular studies identified several instances of inter-specific hybridisation (Rüber et al. 2001; Koblmüller et al. 2007; Nevado et al. 2011). Such examples allowed us to select a case to study the importance of prezygotic, behavioural isolation after, or at the last stages, of the speciation process.

*Ophthalmotilapia* Pellegrin, 1904 species are maternal mouth brooders that occur on the rocky and intermediate (rocky patches separated by sand) shores of Lake Tanganyika. The genus contains four currently accepted valid species: *O. ventralis* (Boulenger 1898), *O. boops* (Boulenger 1901), *O. heterodonta* (Poll and Matthes 1962) and *O. nasuta* (Poll and Matthes 1962) (Hanssens et al. 1999). They are sexually dimorphic maternal mouthbrooders with territorial males that protect a spawning site, and females that aggregate in feeding schools when they are not breeding. *Ophthalmotilapia* males possess egg-shaped lappets at the distal ends of their elongated pelvic fins that are unique among Great Lake cichlids (Poll 1986). These lappets function as egg dummies during the species’ mating behaviour in a similar way as the egg spots on the anal fins of the so-called ‘modern’ haplochromines (sensu Salzburger et al. 2007; Theis et al. 2012). During the mating process, the female deposits the eggs and almost immediately takes them into her mouth. By snapping at the egg dummies, which are situated close to the genital opening of the male, the intake of sperm is thought to be facilitated, increasing the fertilisation rate of the eggs within the female’s mouth (Salzburger et al. 2007).

The four species of *Ophthalmotilapia* have different but partially overlapping, distribution ranges. *Ophthalmotilapia nasuta* is the sole species in the genus with a patchy but lake-wide distribution. The sister species *O. heterodonta* and *O. ventralis* have non-overlapping ranges with the former occurring in the northern half and the latter in the southern third of the Lake. The fourth species, *O. boops* only occurs along a rather limited stretch of Lake Tanganyika’s south-eastern shoreline. There, it prefers sites where large stones are available (Konings 2019). This is the only part of the lake where up to three species of *Ophthalmotilapia* occur in sympatry (Hanssens et al. 1999).

Although specimens of *Ophthalmotilapia* can be easily assigned to one of the valid species (with the possible exception of *O. heterodonta* and *O. ventralis*, see Hanssens et al. 1999), a phylogeographic study discovered gene flow among these species. Nevado et al. (2011) observed that specimens of *O. nasuta* often carried mitochondrial DNA of the other species, whereas the opposite was much less often the case. They suggested that this pattern either has a postzygotic, (*e.g*. by cyto-nuclear incompatibilities that affects mutual crossbreedings differently) or a prezygotic (*e.g*. by an asymmetry in reproductive behaviour that results in a different resistance towards hybridisation) cause. The latter scenario implies that females of all species would occasionally mate with *O. nasuta* males, while *O. nasuta* females would be much less inclined to mate with heterospecific males. It also implies that the female hybrid offspring would backcross into *O. nasuta*. This scenario agrees with the recent description of a successful mating between a female *O. ventralis* and a male *O. nasuta* (Kéver et al. 2018). Reproductive isolation in closely related species of East African cichlids is mostly maintained through prezygotic isolation (Turner et al. 2001). Hence, models that describe speciation in cichlids emphasize the importance of female mate choice in the initial stages of the speciation process (Danley and Kocher 2001).

The Lake Tanganyika cichlids assemblage contains species with profoundly different mating strategies (Ronco et al. 2020). *Ophthalmotilapia* stands out by its extreme sexual dimorphism and female-biased reproductive investment. Although the correlation between reproductive investment, sexual selection and choosiness is well-established, it still remains debated whether choosiness is an evolutionary outcome (sensu Trivers 1972), or rather a determinant of differences between the sexes in parental investment (Thomas and Szekely 2005). Using Lake Tanganyika cichlids, Gonzales-Voyer et al. (2008) showed support for the latter hypothesis. Regardless of the evolutionary mechanism, females should be considered the choosy sex in *Ophthalmotilapia* (sensu Wirtz, 1999). Therefore, if a prezygotic mechanism explains the asymmetric pattern observed in nature (Nevado et al. 2011), it would be caused by differences between females of the different species in accepting matings with heterospecific males. As increased capacity for species recognition leads to increased preference in the choosier sex (Kozak & Boughman, 2009), we predict to see an interspecific difference in female response to conspecific and heterospecific males. As males of the different *Ophthalmotilapia* species have very similar courtship behaviours, in which the few species-specific elements are insufficient to prevent hybridisation (Kéver et al. 2018), females would mainly rely on other cues like colour patterns, body size and pheromones. Although the reproductive behaviour of *Ophthalmotilapia* species is well documented (Haesler et al. 2011; Immler and Taborsky 2009; Kéver et al. 2018), little is known on how *Ophthalmotilapia* species recognize conspecifics and select potential mates.

In this study, we will investigate species recognition, which was, in spite of its shortcomings (Mendelson and Shaw 2012), defined as “a measurable difference in behavioural response toward conspecifics as compared to heterospecifics’’. We will study this by comparing the ability of female *O. nasuta* and *O. ventralis* to distinguish con-from heterospecific males. Often, dichotomous mate choice trails, in which a female can choose between two males, are used for such studies. Such a setup includes an aspect of male competition, whereas we are interested in the response of females in the absence of such behaviour. Additionally, with dichotomous trails one cannot discriminate mate preference from species recognition. Although we expect species recognition to be a good predictor of female mate preference, we did not aim to quantify preference directly. Instead, we choose to investigate differences in species recognition by presenting a set of *O. ventralis* and *O. nasuta* females to males of both species separately and by measuring a range of behavioural variables.

This study described and compared the initial behavioural response of *O. ventralis* and *O. nasuta* females towards males of both species in an aquarium setting. In view of the pattern of asymmetric hybridisation discovered among these species (Nevado et al. 2011), we expected a difference in the capacity for species recognition between females of *O. ventralis* and *O. nasuta*. Specifically, we hypothesized that *O. nasuta* females would be able to differentiate between conspecific and heterospecific males at the initial stages of an encounter. For *O. ventralis* females, however, we expected that this capacity would be less pronounced or absent.

## Methods

### Experimental setting

We performed two independent experiments using females and males of *O. ventralis* and *O. nasuta*. The first focused on the behaviour of focal *O. nasuta* females (ON experiment), the second on that of focal *O. ventralis* females (OV experiment). All individuals were wild caught off the coast of Ulwile Island or nearby Kala, on the mainland (Tanzania). Females were acquired as juveniles and hence had no prior mating experience. They were kept in monospecific aquaria per species until they reached maturity. Hence, all females used had similar ages. Males were older and most had prior mating experience (Kéver et al. 2018). We verified the origin of a random selection of these fishes by sequencing the mitochondrial control region, and by comparing these sequences with the data collected by Nevado et al. (2011). The experiments were performed at the aquarium facilities of the University of Liège. Prior to the onset of the experimental trials, the sex of the specimens was checked by visually inspecting their genital papillae. Female specimens were kept jointly but isolated from males and heterospecific specimens in a separate tank for at least two weeks. This tank was devoid of hiding places, in order to prevent the development of territoriality. During that period, males were kept in monospecific tanks in which they were visually isolated from each other using opaque partitions. We kept all specimens in the same condition for at least two weeks with photoperiod: 12:12 h L:D, water temperature: 26±1°C, carbonate hardness: >8 dKH. Fishes were fed daily ad libitum with ‘Tropical Spirulina forte’ mini-granules.

We used three identical experimental aquaria (88cm*50cm*40cm with water level ca. 40cm), which we divided into two equal parts by a perforated transparent partition (separation wall), through which fishes could not pass, and by an opaque wall (visual barrier) that could be removed (Fig. 1A). A flower pot was placed on each side of the separation to allow the fish to take refuge. We kept the fishes in these aquaria for at least twelve hours before they were used in the experimental trials. During the trial, the visual barrier was removed.

**Figure 1.**
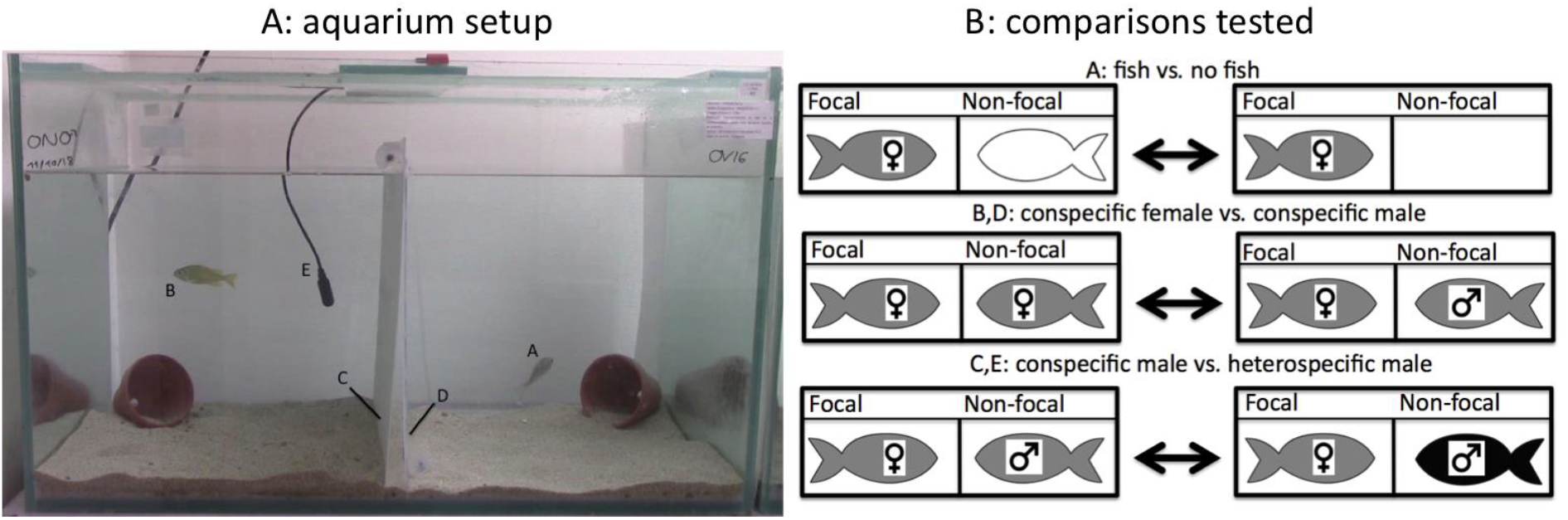
Experimental setup. A: aquarium setup: A focal female of either *O. nasuta* or *O. ventralis* (here *O. nasuta*) (A) was placed in one half of the experimental tank whereas no fish, a conspecific female or a hetero- or conspecific male was placed in the other half (here an *O. ventralis* male) (B). The tank was divided in two by a transparent wall (D) and a visual barrier (C), a hydrophone (E) was placed on the side of the non-focal specimen and an empty flowering pot was placed in both halves of the tank, allowing fishes to take refuge. Video and audio recordings were made 15 minutes prior and 45 minutes after the visual barriers were removed. B: Contrasts tested using permanova: (A): focal females presented with another fish *vs*. with no fish, (B): focal females presented with a conspecific female *vs*. a conspecific male, (C): focal females presented with a conspecific *vs*. a heterospecific male, (D): conspecific females and males presented to a focal female and (E): conspecific and heterospecific males presented to a focal female. Black and grey fishes represent different species, the white fish represents all possible non-focal specimens used.

We recorded the behaviour of focal specimens (*O. nasuta* females or *O. ventralis* females) in four different experimental conditions. They were either exposed to (i) no other specimen (Co), (ii) a conspecific female (CF), (iii) a conspecific male (CM) or (iv) a heterospecific male (HM) (Supplement 1). For each experiment (ON and OV) and for each experimental condition (i to iv), we conducted a minimum of five replicates. We filmed (using a CANON Legria HF R606) the entire aquarium (i.e. focal and non-focal fishes) during one hour: from 15 minutes before to 45 minutes after the visual barrier was removed. Experimenters were only briefly present in the room to remove the visual barrier. As *O. ventralis* males are known to produce weak-pulsed sounds during the inviting behaviour (Kéver et al. 2018), we used a HTI Min-96 hydrophone (−164.4 dB re. 1 V μPa−1; bandwidth 2 Hz and 30 kHz, MS, USA), connected to a Tascam DR-05 recording (TEAC, Wiesbaden, Germany) at a 44.1 kHz sampling rate to record sounds during the whole experiment. The hydrophone was positioned near the separation wall, at half the height of the water column, on the side of the non-focal specimen. At the start of each trial, we switched off the aeration of the tank so that sounds could be recorded. However, these recordings were not analysed, as we detected no communication sounds. After each trial, the focal female was euthanized. Both the focal and the non-focal specimens were weighed. Focal specimens were measured, dissected and the stage of gonad development was scored following Panfili et al. (2006).

We performed a total of 28 ON and 21 OV experimental trials, with a maximum of three trials per day (Supplement 1). However, after the dissections (see below), we observed that six focal *O. nasuta* females from the first set of trials possessed male or ambiguous gonads (Supplement 2). These specimens were referred to as floater males and the recordings for these trials were not analysed. As we suspected that these specimens had changed sex, we photographed the genital papillae of the focal females that were to be used subsequently, two weeks before the onset of the experimental trials. A comparison between papillae of the same individuals after two weeks revealed that some could be identified as males. This either confirmed that a sex change did indeed take place in several specimens, or showed that the examination of genital papillae was insufficient for sex determination in subadult *Ophthalmotilapia*. These specimens were not studied. After each trial, the aquarium was cleaned and the water fully renewed.

### Collection of tracking and qualitative behavioural data

Video files were converted into JPG images using Adapter v2.1.6 (available at https://www.macroplant.com), capturing one frame per second and saving it as an 8-bit, gray-scale JPG file. Images taken within ten seconds before or after an experimenter was performing an action (i.e. removing the wall) were discarded from analyses. We chose to analyse the same number of frames for all trials within the ON and the OV experiment, respectively. For the ON experiment, this resulted in a minimum of 721 and 2186 frames collected before and after the removal of the separation wall, respectively. For the OV experiment, 871 and 2685 frames were available for analyses. Both focal and non-focal specimens were tracked using the ImageJ v1.49 (Schneider et al. 2012) software package.

Given the presence of both light- and dark-coloured backgrounds in the aquarium setting, the set of frames was studied twice. Specimens that were present before a light coloured background were tracked by inverting black and white values whereas specimens present before a dark-coloured background were tracked using non-inverted images. For computational reasons, analyses were performed on subsets of the data containing a maximum of 1,000 frames. For each set of images, a subset of 30 frames was used to create a background using the plug-in ‘*Image stack merger plus*’. Backgrounds were removed using the image calculator and the resulting frames were transformed into black-and-white images using the threshold function with ‘*MaxEntropy*’ as the methodology. Images were adjusted using the ‘*erode*’ and ‘*dilate*’ functions to remove noise and to obtain a better representation of the fishes. The resulting image series was then used for tracking using the plug-in ‘*MTrack2*’, in which tracks were summarized as the x- and y-coordinates of the centroids of the tracked object. The quality of the automated tracking was checked by visually inspecting each of the frames. When the software failed to track a specimen that was clearly present in the final images, coordinates were added manually. Finally, tracks obtained from both datasets, inverted and non-inverted, were combined. When a specimen was recognized by both methods, *e.g*. when the fish was partially before a light- and partially before a dark-coloured background, the average of the coordinates was used. When tracking data was missing, the average value of the coordinates of the previous and the next positions were used. This is justified, as missing data either corresponded to fish that remained stationary for many frames, and could hence not be distinguished from the background, or to fish that hid behind the terracotta flowerpots (Fig. 1). Frames collected before and after the removal of the visual barrier were analysed separately. Coordinates were shifted using the lower- and anterior-most point of the separation wall as the origin, and rotated by setting the anterior water level as a reference for the horizontal plane. Finally, all coordinates were transformed from pixels to centimetres using the dimensions of the aquaria. Tracks were visualized by plotting all individual positions as well as the shift in average position of a specimen before and after the removal of the barrier (Supplement 3).

For each specimen, six tracking parameters were calculated from the coordinates (Table 1). Each parameter was calculated three times: once using coordinates obtained for 721/871 (OV/ON experiment respectively) seconds before (before) the removal of the visual barrier, once using coordinates obtained during 721/871 seconds (after1) after the visual barrier was removed and finally using coordinates obtained during 2186/2685 seconds (after2) after the visual barrier was removed, hence including the after1. Additionally, ten specific behaviours were defined based on Baerends and Baerends-Van Roon (1950). These were encoded and recorded as point events in Boris v. 2.72 open source software (Friard and Gamba 2016) (Table 1). This data was collected during the same three periods: before, after1 and after2. We choose these time periods, as we aimed to study the immediate response following presentation to another fish. Teleosts respond to changing social situations at two scales, by an immediate change in behaviour, and by a modification of neural circuits (Oliveira 2012). The latter is driven by the regulation of immediate early genes (IEG). Following Oliveira (2012), we assumed that the transcription of IEG occurs roughly 15 min (after1), and their translation 45 min (after2) after the stimulus. Behaviours displayed within ten seconds before or after an experimenter was performing an action (removing the wall) were discarded from the analyses.

**Table 1.**
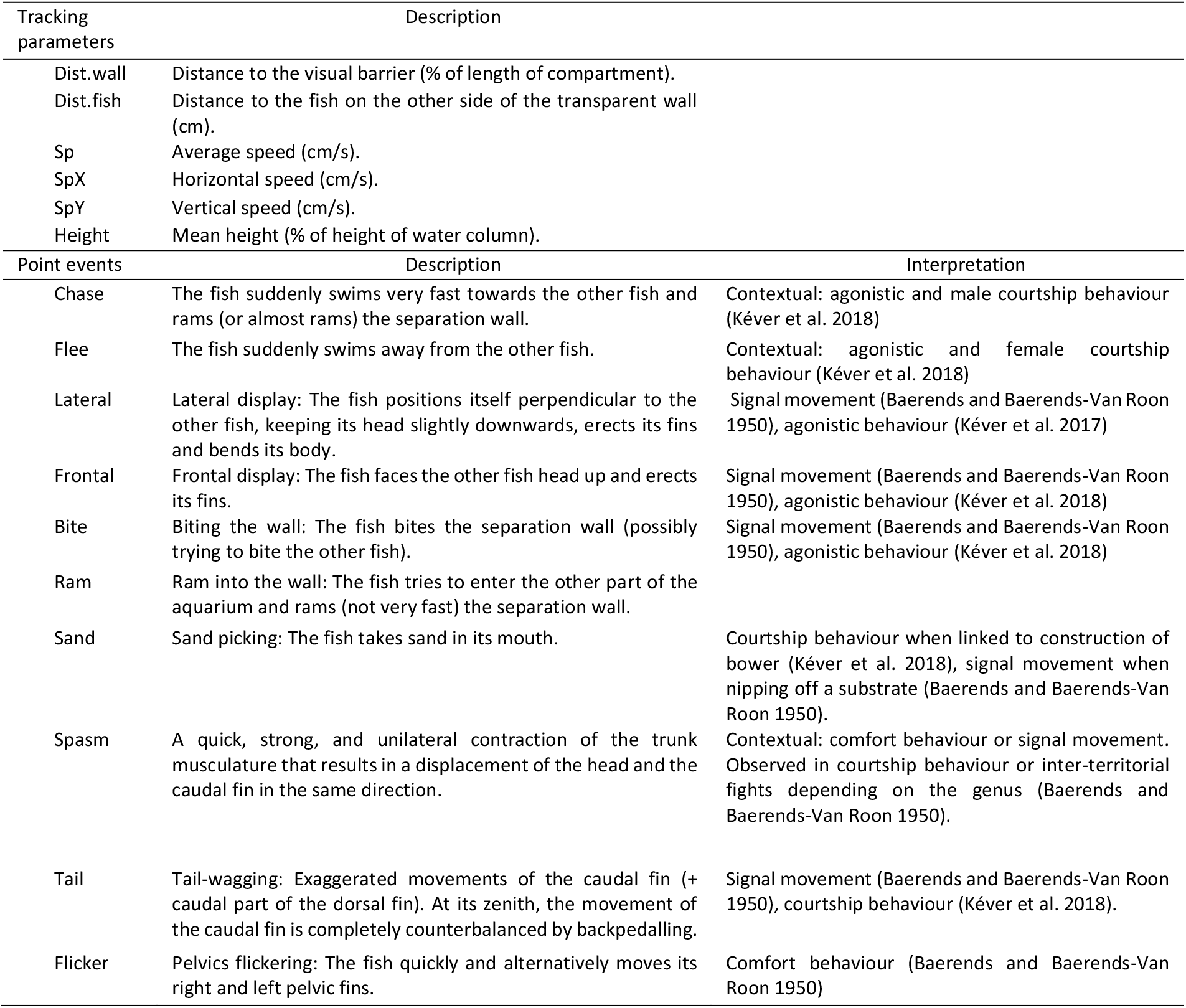
Tracking parameters and point events recorded for the focal and non-focal individuals during the experimental trials. For point events, interpretation of the behaviour was added.

### Statistical analyses

Prior to testing differences in species recognition, we visually explored the combined datasets of tracking parameters and point events. We did this by performing principal component analysis (PCA) and canonical variate analyses (CVA) in Past 3.14 (Hammer et al. 2001). The former allowed for an unbiased visualization of the variation in the data and was performed on the correlation matrices. The latter was conducted to maximize the differentiation between the different groups. Separate analyses were performed for each period of time recorded (before, after1, after2) and for each experiment (ON or OV). Point events that were not recorded during one of these periods were disregarded and missing values (i.e. for Dist.fish in the control condition Co) were treated using mean value imputation. Prior to the analyses, each of the tracking parameters and point events was normalised. This was done for each experiment (OV and ON), and for each of the time periods (‘before’, ‘after1’, ‘after2’ and ‘shift’) separately.

We used permanova to compare the behaviour of both focal and non-focal specimens in five different comparisons (Fig. 1B). With this non-parametric test, we could include all recorded behaviours. This allowed us to measure species recognition via differences in behaviour (sensu Mendelson and Shaw 2012) without defining *a priori* in what variable specimens would differ. In order to reduce the number of comparisons, we restricted ourselves to only biologically relevant contrasts. For focal specimens, we compared the behaviour between (A) females that were presented with another fish *vs*. with no fish, (B) focal females that were presented with a conspecific female *vs*. a conspecific male, and (C) females that were presented with a conspecific *vs*. a heterospecific male. Two additional comparisons were tested for the non-focal individuals. We tested (D) whether conspecific females and males respond differently to a focal female and (E) whether conspecific and heterospecific males respond differently to a focal female. Even though our main goal was to test whether females of *O. nasuta* and *O. ventralis* differed in behavioural response towards conspecific and heterospecific males (C), we tested the four other comparisons as well, following the recommendations of Moran (2003) and Nakagawa (2004).

Tests were performed using non-parametric permanova, using the *pairwise.adonis* function, of the R package *vegan* on the combined data of tracking parameters and point events. This approach was chosen as the conditions for multivariate normality were violated. When permanova revealed significant differences, we verified whether this could be due to dispersion effects (Anderson 2006). For this, in view of the size of the dataset, non-parametric Mann-Whitney U tests were performed on the within group dispersions from the mean, calculated using the *betadisper* function implemented in the R package *vegan* (Oksanen et al. 2017). A difference in dispersion across groups does, however, not contradict a difference in their average values. Hence, the data was visually explored, using PCA, to assess whether dispersion could be present together with a difference in average values. For each comparison revealed significant by permanova, Mann Whitney U non-parametrical tests were performed on each of the variables separately in order to detect which of these caused the difference between the treatments. We choose this non-parametric approach as the assumptions of normality were often not met. When significant, the effect sizes of these variables were estimated using Hedge’s g, which was calculated using the estimationstats.com web application (Ho et al. 2019).

In order to test whether the observed differences in behaviour depended on the visual presence of another specimen, and whether these differences were already visible at the first stages of the encounter, permanova tests were performed on data collected before the removal of the visual separation (before) as well as on data collected over a short (after1) and a long (after2) period of time after this separation was removed. Finally, an additional test was conducted which removed individual variation between the different treatments. For this, behavioural shifts were calculated for each tracking parameter and point event by subtracting the values of the ‘before’ period before from those of the ‘after1’ period (shift). All tests were performed separately for the ON and the OV experiment. As behaviour can be influenced by gonad development and weight of the focal and non-focal specimens, Mann Whitney U tests were performed to check whether these differed between the treatments. Such tests were also performed on the amount of frames in which fishes could not be tracked. All statistical analyses were performed using Past 3.14 (Hammer et al. 2001) and R (R core team 2017).

## Results

We separately analysed two experiments, one focusing on *O. nasuta* and one on *O. ventralis* females (ON and OV experiment). In the ON experiment, two males of *O. ventralis* performed advanced courtship behaviours. After the encounter, these males started to swim in circles, in fast and erratic movements. This was accompanied by tail wagging, generally displayed close to the partition wall. These males often bit the hydrophone and picked up and moved around sand (49 and 45 times within 45 min *vs*. 0 for the other males). One of these two males (*O. ventralis* male presented to ON38) also tried to chase the female (79 times) and presented the egg dummies of its pelvic fins (5 times). This behaviour stopped immediately when the experimenter removed the female fish. During the encounter, these males turned dark grey, to almost black, which was swiftly reversed after the experimental trial. As we designed our experiment to study behavioural response in the absence of courtship behaviour, we removed these outliers from all analyses. All ten point events were observed in at least one of the fishes in the ON experiment, whereas ‘tail’ (*i.e*. tail wagging) was never observed in the OV experiment (Supplement 4).

### Visualization of the behavioural data

We visually explored the data using PCA and CVA to compare the behaviour of all specimens used in each of the two experiments. In the PCAs conducted on the behavioural data collected before the removal of the barrier, values of all females as well as of conspecific males overlapped (Fig. 2A,B), suggesting a highly similar behaviour. However, heterospecific males were (somewhat) separated from all other specimens by their higher values for PC1 (ON experiment) or PC2 (OV experiment). The loadings of the main PCs (Supplement 5.1) suggest that this difference was due to a more active swimming behaviour (Sp, SpX, SpY) higher up in the water column (height) for *O. ventralis* males (ON experiment) and a higher number of point events (ram, sand, bite) performed at the floor of the aquarium (height) for the *O. nasuta* males (OV experiment), prior to their presentation to a heterospecific female. The fact that *O. nasuta* males build true bowers could explain the higher number of point events, especially ‘sand’. Additionally, *O. nasuta* might be less at ease close to the water surface as it generally occurs in deeper waters than *O. ventralis* (Konings 2019). The CVAs also reflected the behavioural differences of heterospecific males (Fig. 3A,B), as they had higher values for the first CVs. The behaviours that contributed strongly to the separating PCs, also contributed to the main CVs (Supplement 5.2).

**Figure 2.**
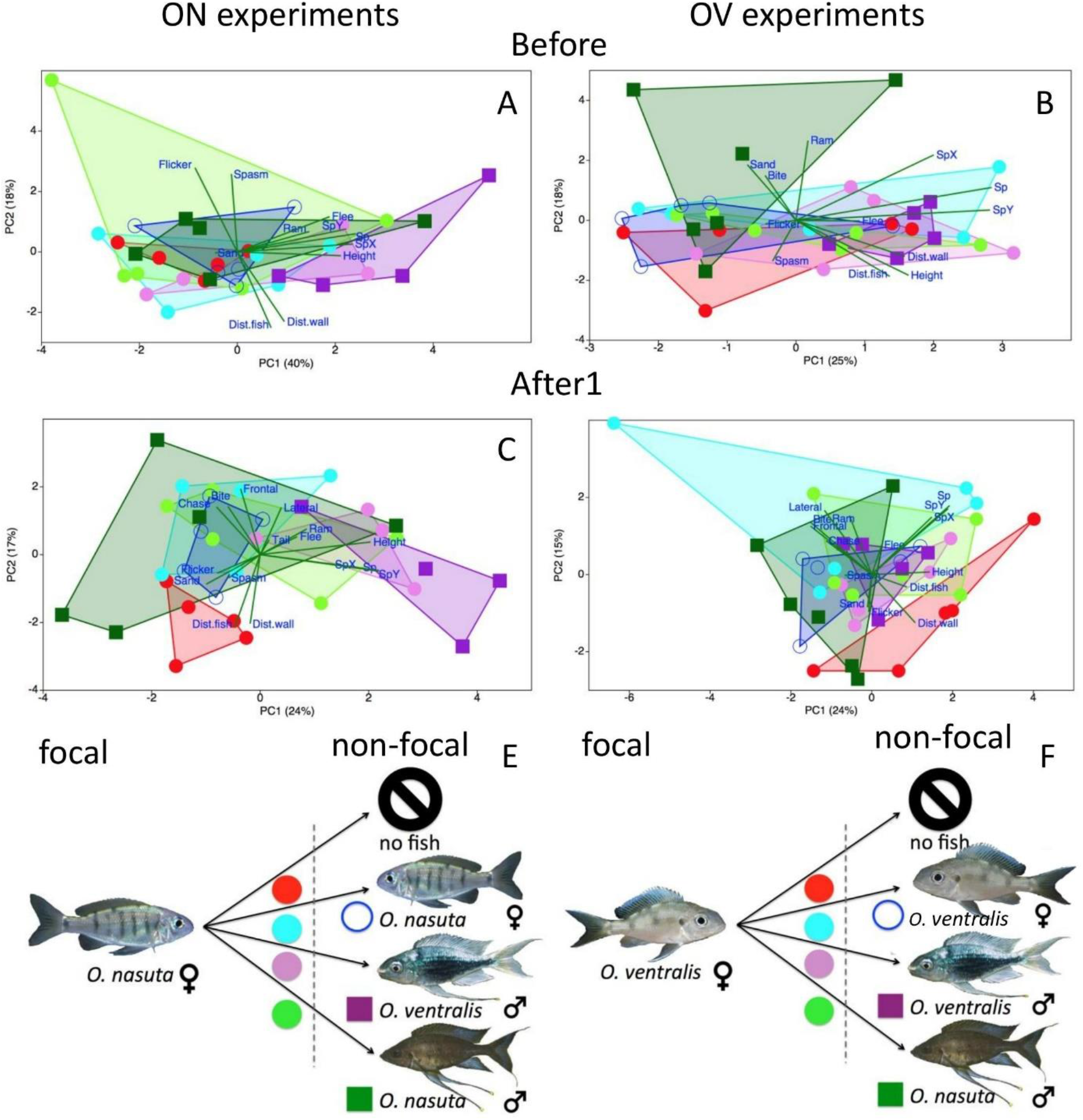
Principal component analyses performed on the behavioural data collected 15 min before (A, B) and 15 min after (C, D) the visual barrier was removed in the ON (*O. nasuta*, left) and the OV (*O. ventralis*, right) experiments. Symbols on the scatter plots for the ON and OV experiment as in E and F, respectively, with full circles denoting focal females presented with no fish (red), a conspecific female (blue) an *O. ventralis* male (purple), and an *O. nasuta* male (green), empty circles denote non-focal conspecific females and full squares *O. ventralis* (purple) and *O. nasuta* (green) males. Explained variances are added to the axes.

**Figure 3.**
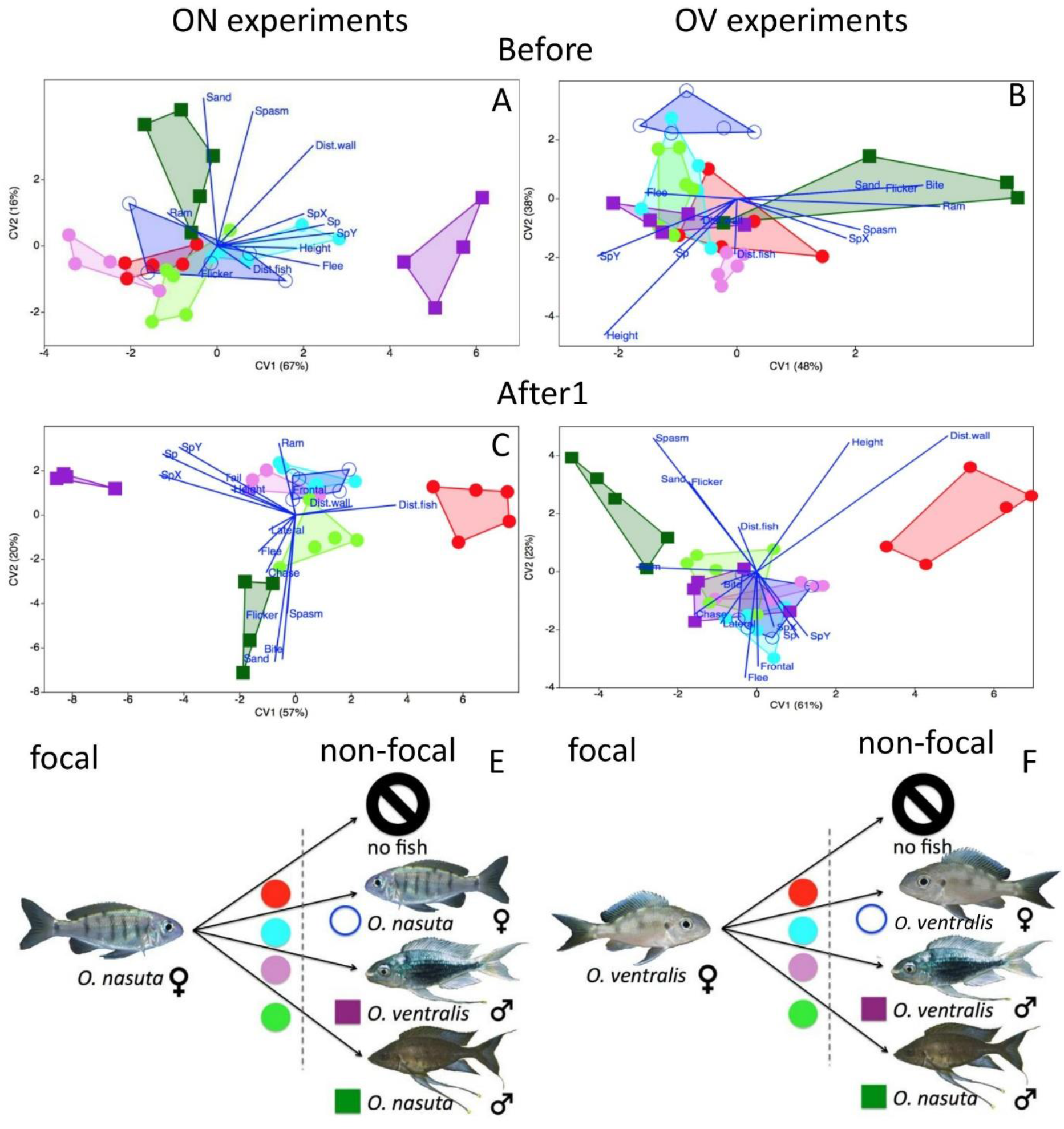
Canonical variate analyses on the behavioural data collected 15 min before and 15 min after the visual barrier was removed in the ON (*O. nasuta*, left) and the OV (*O. ventralis*, right) experiment. Symbols on the scatter plots for the ON and OV experiments as in E and F, respectively, with full circles denoting focal females presented with no fish (red), a conspecific female (blue) an *O. ventralis* male (purple), and an *O. nasuta* male (green), empty circles denote non-focal conspecific females and full squares *O. ventralis* (purple) and *O. nasuta* (green) males. Explained variances are added to the axes.

The PCAs performed on the data collected 15 minutes after the removal of the barrier foremost showed how the focal females that were used as controls behaved differently than the specimens that were presented with another specimen (Fig. 2C,D). For both experiments, this can be explained by control females spending less time closer to the wall (Dist.wall), and performing less agonistic behaviour (ram, lateral, flee). We did not observe any additional separation in the PCA of the OV experiment (Fig. 2D). In the ON experiment, values for (heterospecific) *O. ventralis* males stood out by their high values for PC1, whereas those of (conspecific) *O. nasuta* males had mostly low values for this axis (Fig. 2C). The tracking parameters Sp, SpX, SpY all had high, positive contributions to PC1 (Supplement 5), reflecting that *O. ventralis* males kept swimming actively when the barrier was removed in the ON experiment.

We carried out CVAs on the same datasets (Fig. 3C,D). For both experiments, the control females stood out by their high values for CV1. This could again be explained by their higher values for Dist.Wall. In the ON experiment, (heterospecific) *O. ventralis* males stood out by their low values for CV1, which would be attributed to their more active swimming behaviour (Sp, SpX, SpY). Conspecific *O. nasuta* males stood out by their low values for CV2, which could be due to the higher occurrence of ‘sand’ and ‘bite’ behaviour. Values for *O. nasuta* females that were presented with another fish had more intermediate values for CV1 and CV2. However, values of *O. nasuta* females that were presented to a conspecific male clustered between values of those males and of those of the other females. Similarly, females that were presented to a heterospecific male had values that were intermediate between those of these males and those of the other females (Fig. 3C). This suggests that the behaviour of focal females shares characteristics with the behaviour of the non-focal fishes presented to them. In the CVA of the OV experiment, (heterospecific) *O. nasuta* males stood out by their low values for CV1 and high values for CV2. This was most influenced by the higher occurrence of point behaviours (ram, spasm, sand, flicker). Values of *O. ventralis* males overlapped with those of female specimens that were presented with another fish (Fig. 3D). These patterns remained present when performing similar analyses on the data collected 45 minutes after the removal of the visual barrier (Supplement 6). Plotting the shift in average position before and after the removal of the barrier revealed how almost all specimens moved towards the wall when presented with another specimen. Additionally, this showed that *O. nasuta* specimens, on average, spent more time closer to the bottom whereas *O. ventralis* specimens were more often found higher up in the water column (Supplement 7).

### Behaviour of focal females

We tested whether the behavioural responses of females of *O. nasuta* and *O. ventralis* differed in three different comparisons (Fig. 1B), i.e. when they were presented to (A) another fish *vs*. no fish, (B) a conspecific female *vs*. a conspecific male, and (C) a conspecific *vs*. a heterospecific male. Within each of the comparisons in the ON experiment, we detected no significant difference in the gonad development of the focal females, in their weights, in the weights of the non-focal fishes, and in the percentage of missing frames. The same applies for the OV experiment, although here, non-focal females had lower body weights than conspecific males. Prior to the removal of the visual barrier, no significant difference in behaviour was recorded for focal females from the different treatments in both experiments, and for all three comparisons (Table 2).

**Table 2.** PERMANOVA performed on the behavioural parameters of the ON (*O. nasuta*) and OV (*O. ventralis*) experiment. Tests were performed on the data collected during 15 minutes before (B), 15 minutes after (A1) and 45 minutes after (A2) the removal of the opaque wall as well as on behavioural shifts (S) calculated by subtracting data recorded during 15 minutes before from that of 15 min after the removal of the wall (A1-B). We tested five comparisons (Fig. 1B): by comparing the behaviour of focal females that were presented with (A) another fish *vs*. with nothing, (B) with a conspecific female *vs*. a conspecific male, and (C) with a conspecific *vs*. a heterospecific male. We further compared (D) the behaviour of non-focal conspecific males vs. females, and (E) conspecific *vs*. heterospecific males. Behaviours that were neither recorded before or after the removal of the wall, ‘tail’ and ‘sand’ for the ON, and ‘flicker’ and ‘sand’ for the OV experiment, were excluded. For the first comparison, Dist.fish was excluded, as it could not be calculated. Values in bold are significant at the 0.05 level. For these comparisons, ° and † denote that the assumption of equal dispersion was violated at the 0.05 and 0.01 levels.

|  |  | A | B | C | D | E |
| --- | --- | --- | --- | --- | --- | --- |
| ON | B | F:0.31; p: 0.789 | F:0.95; p: 0.423 | F:0.91; p: 0.474 | F:1.17; p: 0.339 | F:1.15; p: 0.24 |
|  | A1 | **F:7.28; p: 0.012 †** | F:0.42; p: 0.768 | **F:11.33; p: 0.016 °** | F:0.45; p: 0.651 | F:0.55; p: 0.73 |
|  | A2 | **F:4.71; p: 0.027 †** | F:0.49; p: 0.553 | **F:8.13; p: 0.040** | F:0.66; p: 0.545 | F:0.85; p: 0.47 |
|  | S | **F:8.40; p: 0.006 †** | F:1.03; p: 0.416 | **F:8.75; p: 0.018 †** | F:1.18; p: 0.332 | F:0.25; p: 0.93 |
| OV | B | F:0.23; p: 0.762 | F:1.62; p: 0.203 | F:0.85; p: 0.387 | F:3.19; p: 0.061 | **F:5.13; p: 0.013** |
|  | A1 | **F:8.00; p: 0.005** | F:0.91; p: 0.386 | F:0.40; p: 0.546 | F:0.28; p: 0.633 | F:0.48; p: 0.524 |
|  | A2 | **F:4.28; p: 0.026** | F:0.78; p: 0.438 | F:0.48; p: 0.610 | F:0.09; p: 0.837 | F:0.49; p: 0.525 |
|  | S | **F:7.61; p: 0.004** | F:0.89; p: 0.418 | F:0.87; p: 0.407 | F:0.42; p: 0.679 | F:0.73; p: 0.447 |

For the three comparisons, and for both experiments (ON and OV), we performed permanova on the behavioural data recorded during the 15 minutes after the removal of the visual barrier (Table 2). For both the OV and the ON experiment, this revealed a significant difference in the behaviour of focal females that were not presented to another fish (controls) and focal females that were presented with another fish (comparison A). For the ON experiment, we could not exclude that these differences stem from dispersion effects. However, these dispersion effects can be explained by the (much) lower variation in values for control females compared to those of other focal females. As visualisation of the data showed a clear separation of both groups, we can assume that a difference in dispersion is present together with a difference in average values (Fig. 2C). Mann-Whitney U tests revealed that controls differed from other focal females by their higher values for the variable Dist.wall (ON: 46.3 +-18.2 *vs*. 20.1 +-8.8, p=0.005, g=1.32; OV: 55.2 +-13.3 *vs*. 20.8 +-9.7, p=0.002, g=1.13) and their lower values for ‘ram’ (ON: 8.6+-12.1 *vs*. 74.5+-54.0, p=0.005, g=-2.14; OV: 8.0 +-11.9 *vs*. 48.8 +-38.4, p=0.014, g=-3.12).

Unexpectedly, in both experiments, we did not observe a difference in behaviour between focal females that were presented with a conspecific female or a conspecific male (comparison B). However, when comparing the behaviour of focal females presented with conspecific and heterospecific males (comparison C), a difference became evident between the ON and the OV experiment. In support of our hypothesis, females of *O. nasuta* responded differently towards conspecific and heterospecific males, whereas females of *O. ventralis* did not (Table 2). Mann-Whitney U tests revealed that this was due to the lower number of observed ‘ram’ behaviours (43.8+-17.9 *vs*. 138+-52.9; p=0.04, g=2.25) in *O. nasuta* females that were presented to *O. nasuta* males compared to those presented to *O. ventralis* males. Additionally, *O. nasuta* females never performed a ‘flee’ behaviour when presented to an *O. ventralis* male, whereas this was observed in all but one of the *O. nasuta* females presented to a *O. nasuta* male (0 *vs*. 2.6+-2.3; p=0.04, g=-1.33). We obtained highly similar results when analysing the data collected 45 minutes after the visual barrier was removed, or when analysing the shift data (Table 2).

### Behaviour of non-focal specimens

We tested whether the behavioural response of conspecific females and males differed when presented to a focal female (D) and whether the behavioural response of conspecific and heterospecific males differed when presented to a focal female (E). Unexpectedly, permanova revealed a difference in the behaviour of *O. nasuta* and *O. ventralis* males in the OV experiment, prior to the removal of the barrier. Mann-Whitney U tests revealed that this was due to the higher average vertical swimming speed of *O. ventralis* males compared to *O. nasuta* males (SpY 1.7+-0.5 *vs*. 0.7+-0.4, p= 0.014, g=2.05). None of the other comparisons was shown to be significantly different, neither before, nor after the removal of the barrier (Table 2).

## Discussion

### Summary

We tested the initial response of *O. nasuta* and *O. ventralis* females towards conspecific and heterospecific males. In support of our hypothesis, *O. nasuta* females differentiated between conspecific and heterospecific males, whereas *O. ventralis* females did not. Visualisation of the data revealed that *O. nasuta* females mirrored the behaviour of the males to which they were presented. We also presented females of both species with a conspecific female or with nothing (control). Although females always responded to the presence of another fish, their behaviour did not differ when presented with conspecific males and females. Comparisons of non-focal specimens did not reveal any differences in behaviour after presentation to a focal female. However, before the removal of the wall, males of *O. ventralis* and *O. nasuta* behaved differently in the OV experiment.

### Responses of *Ophthalmotilapia* females

Nevado et al. (2011) discovered signatures of unidirectional hybridisation in *Ophthalmotilapia*, which could either be explained by cyto-nuclear incompatibilities, or by asymmetric mate choice. The latter explanation implies that *O. nasuta* females would discriminate more strongly against heterospecific males, than females of *O. ventralis*. This is supported by our experiments.

As we did not present focal females with heterospecific females, we cannot say that the observed species recognition in *O. nasuta* females was due to a different response towards males, or towards all specimens of the other species. However, as females in *Ophthalmotilapia* are non-territorial and therefore often encounter heterospecific congeners, we expect that the female response is specific to heterospecific males. Unexpectedly, females of both species behaved similarly towards conspecific males and females. Hence, we cannot say whether we observed sexually motivated behaviour, or just the routine behaviour of a (isolated) female that encounters a conspecific individual. In the wild, non-breeding females of both species aggregate in large feeding groups (Konings 2019). Hence, being isolated for 12 hours, as was the case prior to the start of the experiment, represents an unnatural situation for *Ophthalmotilapia* females. It would, therefore, not be unlikely if *Ophthalmotilapia* females are behaviourally hardwired to reunite immediately with conspecifics, regardless of whether these are female or male.

Females of *O. nasuta* only performed the ‘flee’ behaviour towards conspecific males and displayed the ‘ram’ behaviour less frequently. Although we found no evidence that this behaviour is sexually motivated, male chasing and female fleeing (i.e. ‘flee’) form the first steps in the mating process of *Ophthalmotilapia* (Kéver et al. 2018). The ram behaviour, on the other hand, was seen in all experimental trials in which a focal female was presented to another fish. Indeed, the main difference in behaviour between focal females of both species that were, or were not, presented with another fish was the amount of time spent close to the wall, and in the display of the ‘ram’ behaviour.

We discovered that several of the presumed females that we planned to use in the experiments had been, or changed into, males. This was also observed for one of the non-focal females in the ON experiment, which was kept isolated from males after the experiment (Supplement 2). Although there are several reports of sex changes occurring in cichlids (Peters 1975; Naish and Ribbink 1990), evidence hereof remained, until now, limited. As no reproductions were recorded when these specimens still had a female morphology, we cannot rule out that they were males that delayed the development of the conspicuous male colouration and the elongated pelvic fins. Hence, we did not provide solid evidence of sex change in *Ophthalmotilapia*.

### Interpretation

Although we uncovered a significant difference in the behaviour of *O. nasuta* females that were presented with a conspecific and a heterospecific male, permanova did not reveal a significant interspecific difference in the behaviour of the males. This could indicate that *O. nasuta* females interpreted behaviour differently when displayed by *O. nasuta* or by *O. ventralis* males. Such species- or sex-dependent interpretation of behaviour is known for several cichlid species, in which territorial males present themselves identically towards both visiting females and intruding males (Baerends and Baerends-Van Roon 1950). Visual exploration of the data, however (Fig. 3C, Supplement 6), revealed a potential difference in male behaviour, which was mirrored by female response. As female *O. ventralis* did not appear to differentiate between conspecific and heterospecific males, one could ask why hybridisation is not even more prevalent. However, we only examined the very first stage in a potential mating process, so other differences that are present in the mating behaviour between both species could be responsible for this. For example, *O. ventralis* males display a specific late mating behaviour, called ‘invite’, which *O. nasuta* males never display (Kéver et al. 2018). Additionally, hybrids might have a lower fitness. In order to reach mitochrondrial introgression, female hybrids would also need to mate with *O. nasuta* males. This is not unlikely given that, in cichlids, female mate choice is influenced, via imprinting, by the maternal phenotype (Verzijden and ten Cate 2007; Verzijden et al. 2008).

### The role of males

Asymmetric propensities towards hybridisation are expected in the intermediate stages of reproductive isolation (Arnold et al. 1996). This has been observed in a variety of animal taxa including lungless salamanders (Verrell 1990), spadefoot toads (Pfennig 2007), swordtails (Crapon de Caprona and Ryan 1990), pupfishes (Strecker and Kodric-Brown 1999), and several cichlids (Egger et al. 2008; Nevado et al. 2011). Although most examples are related to female mate choice, these patterns can also be caused by asymmetries in male choosiness (Svensson et al. 2007). Although male mate choice is common in fishes (Wong and Jennions 2003; Werner and Lotem 2003), we choose to focus on the role of females (see introduction, Seehausen et al. 2008; Sefc et al. 2017). However, the mode of fertilization in *Ophthalmotilapia* could also have an influence on male choosiness. Haesler et al. (2011) studied the reproductive behaviour of *O. ventralis*, but it can be assumed that the behaviour of its congeners is highly similar. In *O. ventralis*, a ripe female will visit the territories of several males, either to spawn, or just to collect additional ejaculates. Subsequently, sperm competition will take place within her mouth, resulting in clutches with multiple sires (Haesler et al. 2011). Given that this dilutes the effect of a ‘wrong’ choice, a female can afford to be less choosy. Differences in both male and female courtship effort towards genetically distant or similar mates have been documented in another mouth brooding cichlid: *Tropheus* Boulenger, 1898 (Zoppoth et al. 2013). However, *Tropheus* species are sexually monomorphic and both sexes are territorial. Additionally, *Tropheus* males invest significantly more in raising the clutch, by providing the female access to their feeding territories. As males of *Ophthalmotilapia* do not share their resources, we can expect these males to be less choosy than those of *Tropheus*. Additionally, a substantial role of male mate choice is not supported by our data, as we did not observe a difference in behaviour between non-focal males of *O. nasuta* and *O. ventralis* when presented with females of the two species. It should be noted, however, that two *O. ventralis* males that displayed mating behaviour towards *O. nasuta* females were excluded from the analyses.

Whereas our experiments only revealed the capacity for species recognition in females of *O. nasuta*, we cannot conclude that males cannot distinguish between females of the two species. Whereas the males of *O. nasuta* and *O. ventralis* behaved differently when a visual barrier was present, no significant difference was found after its removal. This could imply that males of the two species behave in a similar way when presented with a conspecific or a heterospecific female. However, an alternative explanation would be that males recognise conspecific and heterospecifics, and use this knowledge to court females using a repertoire appropriate to the species. Although this was found in sister species pairs of freshwater sticklebacks (*Gasterosteus* spp. L. 1758) (Kozak et al. 2009), our experimental design did not allow us to test this in *Ophthalmotilapia*.

We cannot exclude that morphological, physiological and behavioural features that distinguish *O. nasuta* males from males of congeners could have caused the asymmetric pattern of introgression. Foremost, *O. nasuta* males become larger and possess longer pelvic fins. This feature could render them more attractive as *O. ventralis* females have a preference towards males with strongly elongated pelvic fins (Haesler et al. 2011). As a change in the feature associated with attractiveness can alter species recognition in the mating process (Phelps et al. 2006), the extra-long pelvic fins of *O. nasuta* males could serve as a super-natural stimulus (sensu Tinbergen 1948). Additionally, even though Haesler et al. (2011) found no correlation between female choice and male body length in *O. ventralis*, they did observe that larger males outcompeted their rivals in sperm competition within the females’ mouth. Additionally, sperm of *O. nasuta* remains viable for a significantly longer amount of time than that of *O. ventralis* (Morita et al. 2014). Lastly, *O. nasuta* males construct true bowers (elaborate, crater-shaped sand mounts), whereas the nests of males of the other species of *Ophthalmotilapia* only consist of a small area of cleaned rock, or of a small pit in the sand (Konings 2019).

### The importance of visual cues

Although animals can use multiple kinds of cues to assess the quality of a potential mate, their final assessment depends on the overall information available. This is exemplified by female mate choice in the allopatric swordtail species *Xiphophorus nigrensis* Rosen 1960 and *X. pygmaeus* Hubbs and Gordon 1943. Here, mating preferences differed depending on whether visual, olfactory or a combination of both cues were available (Crapon de Caprona and Ryan 1990). Different responses to visual and olfactory cues were also shown for females of sympatric *Cyprinodon* Lacipède, 1803 pupfish species from Lake Chichancanab (Mexico). Here, different degrees of asymmetric discrimination of males were observed depending on whether females had access to visual or olfactory information (Strecker and Kodric-Brown 1999). In species-rich systems and in species that form leks, such as *Ophthalmotilapia* spp., females must be able to rapidly assess the quality of a potential mate (Barlow 2002). *Ophthalmotilapia* females could use the morphological characteristics, conspicuous bowers and/or stereotypical displays of males to distinguish them from sympatric congeners. However, even though multiple cues can be involved, mate choice decisions in radiations are often based on just a small amount of (combinations of) these traits (Hohenlohe and Arnold 2010).

The separation wall used in our experiments contained holes that allowed for the exchange of water between both compartments. Hence, besides visual clues, the fishes most likely also received olfactory and acoustic information. Although visual cues were suggested to be the primary factor in species-isolating, female mate choice in other cichlids (Jordan et al. 2003; Kidd et al. 2006), we cannot rule out the importance of other types of information. Studies have shown that olfactory (Blais et al. 2009; Plenderleith et al. 2005), acoustic (Nelissen 1978; Amorim et al. 2004; Kéver et al. 2018) and behavioural (Barlow 2002) information can also influence the mating process. Although Seehausen and van Alphen (1998) showed a certain hierarchy of information, where other cues are taken into account when visual information is absent or masked, other experiments showed that female cichlids are more likely to select the right male when both olfactory and visual cues are available (Plenderleith et al. 2005; Blais et al. 2009). When visual information suffices for mate recognition, the behaviour throughout the mating process, i.e. potentially leading toward spawning, does not need to diverge between closely-related species (Barlow 2002). This may explain why spawning behaviour of *Ophthalmotilapia* is remarkably similar across the genus (Kéver et al. 2018) and why differently-coloured, sympatric mbuna cichlids from Lake Malawi have identical courtship behaviours (McElroy and Kornfeld 1990).

### Ecological reasons for asymmetric hybridisation

Although they can be found in sympatry, *O. ventralis* is more associated with the rocky shores of Lake Tanganyika, whereas *O. nasuta* has a wider ecological range. At rocky shores, *O. ventralis* can be one of the most abundant cichlid species (Sturmbauer et al. 2008). Hence, for an *O. ventralis* female, a random encounter with another *Ophthalmotilapia* male is much more likely to result in a conspecific than a heterospecific encounter. In contrast, for an *O. nasuta* female venturing into the preferred *O. ventralis* habitat, a conspecific encounter would be less often the case. Therefore, the ability to discriminate between conspecific and heterospecifics would be less important for females of *O. ventralis* than for those of *O. nasuta*. A similar interpretation was given to explain asymmetries in female discrimination of sympatric *Cyprinodon* species, where the species with the highest abundance had the lowest choosiness (Strecker and Kodric-Brown 1999). Although a species’ propensity for discrimination could be a consequence of its distribution range, the opposite could also hold. Species that are better in recognizing conspecifics are more likely to maintain the integrity of their gene pool. Hence, they could be better in colonising habitats that have already been occupied by related species. Finally, we showed that substantial behavioural differences can be observed between closely-related species. This should be a warning to be cautious when assuming similarities in the behaviour of certain (model) organisms and related taxa.

## Data, script and code availability

The data and script are deposited on Zenodo under: https://doi.org/10.5281/zenodo.5534485.

## Supplementary material

Supplementary material is available on line under: https://doi.org/10.1101/2021.08.07.455508.

## Ethical note

All specimens were obtained from a certified commercial supplier (Cichlidenstadl, Alerheim, Germany). Specimens were housed in the aquarium facilities of the University of Liège. Experimental procedures were performed in accordance with Belgian law, and approved by the University of Liège Institutional Animal Care and Use Committee (protocol #1759) in accordance with the regulations of the ethical committee of the University of Liège. All manipulations were performed by a FELASA-certified experimenter.

## Acknowledgements

We are grateful to Siegfried Loose for his advice on the rearing of *Ophthalmotilapia* and to Nedim Tüzün for his advice on the tracking software. We thank Ad Konings (Cichlid press) to provide us with pictures of *Ophthalmotilapia*. This study is part of the GENBAS project, which was funded by the BELSPO (Belgian Science Policy) BRAIN-be program. Version 3 of this preprint has been peer-reviewed and recommended by Peer Community In Zoology (https://doi.org/10.24072/pci.zool.100010).

## Supplementary material

Supplementary material is available on line under: https://doi.org/10.1101/2021.08.07.455508.

Supplement 1. Summary of the experiments.

Supplement 2. Indications for sex change observed in *O. nasuta* females. Top row: gonads of some of the specimens that were female in external phenotype but had male or ambiguous gonads (ON33, 35, 26, 43). Vertical left: female gonads in developmental stage 3 (ON34), 4 (ON24) and 5 (ON37) respectively (Panfili et al. 2006). Bottom: two presumed non-focal *O. nasuta* females used in the ON experiment, one of which could have undergone transition after the experiments, and horizontal right, the ventral area of the same specimens, with A: anal pore and UG: urogentital pore.

Supplement 3. Visualization of tracking data. The position of each fish is plotted for each second in which specimens were tracked with the positions recorded before and after the removal of the opaque wall (grey) coloured differently. Ellipses denote the area in which 90% of tracks are situated, large dots denote the average positions before and after the removal of the barrier and arrows shows the change in mean position. The separation wall is visualized as a meshed partition. The average speed before (v0) and after (v1) the removal of the barrier is plotted for each tracked specimen with data for focal specimens given in bold. Abbreviations (ON: *O. nasuta*, OV: *O. ventralis*, F: female, M; male)

Supplement 4. Summary of the data.

Supplement 5. Loadings and variance of the main axes of the PCAs and CVAs conducted in the study.

Supplement 6. Principal component analyses and Canonical variate analyses performed on the behaviours recorded 45 min after the visual barrier was removed of the ON (*O. nasuta*, left) and the OV (*O. ventralis*, right) experiment. Symbols on the scatter plots for the ON and OV experiment as in E and F, respectively, with full circles denoting focal females presented with no fish (red), a conspecific female (blue) an *O. ventralis* male (purple), and an *O. nasuta* male (green), empty circles denote non-focal conspecific females and full squares *O. ventralis* (purple) and *O. nasuta* (green) males. Explained variances are added to the axes.

Supplement 7. Shift in average position of the specimens analysed 15min before and 15min after the removal of the separation wall for the ON (above) and OV (below) experiment. Focal specimens are all visualised on the left, and non-focal specimens on the right. Dimensions in cm, with the vertical bar representing the separation wall. Dashed arrows represent individual fishes, bold arrows the average per treatment. Colours, for focal females (ON and OV) presented with no fish (red), a conspecific female (turquoise) an *O. ventralis* male (pink), and an *O. nasuta* male (light green), and non-focal specimens (right): conspecific females (blue), *O. ventralis* males (purple) and *O. nasuta* males (green).

## References

Amorim MCP, Knight ME, Stratoudakis Y, Turner GF (2004) Differences in sounds made by courting males of three closely related Lake Malawi cichlids. J Fish Biol. 65(5):1358–1371. https://doi.org/10.1111/j.0022-1112.2004.00535.x

Anderson MJ (2006) Distance-based tests for homogeneity of multivariate dispersions. Biometrics. 62(1):245–253. https://doi.org/10.1111/j.1541-0420.2005.00440.x

Arnold SJ, Verrell PA, Tilley SG (1996) The evolution of asymmetry in sexual isolation: A model and a test case. Evolution 50:1024–1033. https://doi.org/10.1111/j.1558-5646.1996.tb02343.x

Barlow GW (2002) How behavioural studies contribute to the species problem: a piscine perspective. Fish Fish. 3(3):197–212. https://doi.org/10.1046/j.1467-2979.2002.00083.x

Baerends GP, Baerends-Van Roon JM (1950) An introduction to the study of the ethology of the cichlid fishes, Leiden, The Netherlands, E. J. Brill.

Blais J, Plenderleith M, Rico C, Taylor MI, Seehausen O, van Oosterhout C, Turner GF (2009) Assortative mating among Lake Malawi cichlid fish populations is not simply predictable from male nuptial colour. BMC Evol Biol. 9(53):53. https://doi.org/10.1186/1471-2148-9-53

Coyne JA, Orr HA (2004) Speciation. Sunderland, MA: Sinauer Associates.

Crapon de Caprona MD, Ryan MJ (1990) Conspecific mate recognition in swordtails, *Xiphophorus nigrensis* and X. pygmaeus (Poeciliidae): olfactory and visual cues. Anim Behav. 39(2):290–296. https://doi.org/10.1016/S0003-3472(05)80873-5

Danley PD, Kocher TD (2001) Speciation in rapidly diverging systems: lessons from Lake Malawi. Mol Ecol. 10:1075–1086. https://doi.org/10.1046/j.1365-294X.2001.01283.x

Egger B, Obermüller B, Eigner E, Sturmbauer C, Sefc KM (2008) Assortative mating preferences between colour morphs of the endemic Lake Tanganyika cichlid genus *Tropheus*. Hydrobiologia. 615:37–48. https://doi.org/10.1007/978-1-4020-9582-5_3

Friard O, Gamba M (2016) BORIS: A Free, Versatile Open-Source Event-Logging Software for Video / Audio Coding and Live Observations. Methods Ecol Evol. 7(11):1–6. https://doi.org/10.1111/2041-210X.12584

Gonzalez-Voyer A, Fitzpatrick JL, Kolm N (2008) Sexual selection determines parental care patterns in cichlid fishes. Evolution 62(8): 2015–2026. https://doi.org/10.1111/j.1558-5646.2008.00426.x

Haesler MP, Lindeyer CM, Otti O, Bonfils D, Heg D, Taborsky M (2011) Female mouthbrooders in control of pre- and postmating sexual selection. Behav Ecol. 22(5):1033–1041. https://doi.org/10.1093/beheco/arr087

Hammer Ø, Harper DA, Ryan PD (2001) PAST: Paleontological statistics software package for education and data analysis. Palaeontologia Electronica. 4:9pp.

Hanssens M, Snoeks J, Verheyen E (1999) A Morphometric Revision of the Genus *Ophthalmotilapia*(Teleostei, Cichlidae) from Lake Tanganyika (East Africa). Zool J Linn Soc-Lon. 125(4):487–512. https://doi.org/10.1111/j.1096-3642.1999.tb00602.x

Ho J, Tumkaya T, Aryal S, Choi H, Claridge-Chang A (2019) Moving beyond P values: data analysis with estimation graphics. Nat Met. 16(7):565–566. https://doi.org/10.1038/s41592-019-0470-3

Hohenlohe PA, Arnold SJ (2010) Dimensionality of mate choice, sexual isolation, and speciation. PNAS. 107(38):16583–16588. https://doi.org/10.1073/pnas.1003537107

Immler S, Taborsky M (2009) Sequential polyandry affords post-mating sexual selection in the mouths of cichlid females. Behav Ecol Sociobiol. 63(8):1219–1230. https://doi.org/10.1007/s00265-009-0744-3

Jiggins CD, Salazar C, Linares M, Mavarez J (2008) Hybrid trait speciation and *Heliconius* butterflies. Philos T R Soc B. 363(1506):3047–3054. https://doi.org/10.1098/rstb.2008.0065

Jordan R, Kellogg K, Juanes F, Stauffer J (2003) Evaluation of female mate choice cues in a group of Lake Malawi mbuna (Cichlidae). Copeia. 2003(1):181–186. https://doi.org/10.1643/0045-8511(2003)003[0181:EOFMCC]2.0.CO;2

Kéver L, Parmentier E, Derycke S, Verheyen E, Snoeks J, Van Steenberge M, Poncin P (2018) Limited possibilities for prezygotic barriers in the reproductive behaviour of sympatric *Ophthalmotilapia* species (Teleostei, Cichlidae). Zoology. 126:71–81. https://doi.org/10.1016/j.zool.2017.12.001

Kidd MR, Danley PD, Kocher TD (2006) A direct assay of female choice in cichlids: all the eggs in one basket. J Fish Biol. 68(2):373–384. https://doi.org/10.1111/j.0022-1112.2006.00896.x

Kocher TD (2004) Adaptive evolution and explosive speciation: the cichlid fish model. Nat Rev Genet. 5(4):288–298. https://doi.org/10.1038/nrg1316

Koblmüller S, Duftner N, Sefc KM, Aibara M, Stipacek M, Blanc M, Egger B, Sturmbauer C (2007) Reticulate phylogeny of gastropod-shell-breeding cichlids from Lake Tanganyika - the result of repeated introgressive hybridisation. BMC Evol Biol. 7:7. https://doi.org/10.1186/1471-2148-7-7

Konings A (2019) Tanganyika cichlids in their natural habitat. 4th edition. El Paso, TX: Cichlid press.

Kozak GM, Boughman JW (2009) Learned conspecific mate preference in a species pair of sticklebacks. Behav Ecol. 20:1282–1288. https://doi.org/10.1093/beheco/arp134

Kozak GM, Reisland M, Boughmann JW (2009) Sex differences in mate recognition and conspecific preference in species with mutual mate choice. Evolution. 63:353–365. https://doi.org/10.1111/j.1558-5646.2008.00564.x

Luttbeg B, Towner MC, Wandesforde-Smith A, Mangel M, Foster SA (2001) State-dependent mate-assessment and mate-selection behavior in female threespine sticklebacks (*Gasterosteus aculeatus*, Gasterosteiformes: Gasterosteidae). Ethology. 107(6):545–558. https://doi.org/10.1046/j.1439-0310.2001.00694.x

Mayr E (1982) The growth of biological thought: Diversity, evolution and inheritance. Cambridge, MA: Harvard University Belknap Press.

Marques DA, Meier JI, Seehausen O (2019) A combinatorial view on speciation and adaptive radiation. Trends Ecol Evol. 34(6):531–544. https://doi.org/10.1016/j.tree.2019.02.008

Mendelson TC, Shaw KL (2012) The (mis)concept of species recognition. Trends Ecol Evol. 27(8):421–427. https://doi.org/10.1016/j.tree.2012.04.001

McElroy DM, Kornfeld I (1990) Sexual selection, reproductive behaviour, and speciation in the mbuna species fock of Lake Nyasa (Pisces: Cichlidae). Environ Biol Fish. 28:273–284.

Moran MD (2003) Arguments for rejecting the sequential Bonferroni in ecological studies. Oicos. 100(2):403–405. https://doi.org/10.1034/j.1600-0706.2003.12010.x

Morita M, Awata S, Yorifuji M, Ota K, Kohda M, Ochi H (2014) Bower-building behaviour is associated with increased sperm longevity in Tanganyikan cichlids. J Evolution Biol. 27(12):2629–2643. https://doi.org/10.1111/jeb.12522

Naish KA, Ribbink AJ (1990) A preliminary investigation of sex change in *Pseudotropheus lombardoi* (Pisces: Cichlidae). Environ Biol Fish. 28:285–294.

Nakagawa S (2004) A farewell to Bonferroni: the problem if low statistical power and publication bias. Behav. Ecol. 15:1044–1045. https://doi.org/10.1093/beheco/arh107

Nelissen MHJ (1978) Sound production by some Tanganyikan cichlid fishes and a hypothesis for the evolution of their communication mechanisms. Behaviour. 64(1-2):137–147. https://doi.org/10.1163/156853978X00477

Nevado B, Fazalova V, Backeljau T, Hanssens M, Verheyen E (2011) Repeated unidirectional introgression of nuclear and mitochondrial DNA between four congeneric Tanganyikan cichlids. Mol Biol Evol. 28(8):2253–2267. https://doi.org/10.1093/molbev/msr043

Oksanen JF, Blanchet G, Friendly M, Kindt R, Legendre P, McGlinn D, Minchin PR, O’Hara RB, Simpson GL, Solymos PM, Stevens HH, Szoecs E, Wagner H (2017) vegan: Community Ecology Package. R package version 2.4-5.

Panfili J, Thior D, Ecoutin J-M, Ndiaye P, Albaret J-J (2006) Influence of salinity on the size at maturity for fish species reproducing in contrasting West African estuaries. J Fish Biol. 69(1):95–113. https://doi.org/10.1111/j.1095-8649.2006.01069.x

Peters HM (1975) Hermaphroditism in cichlid fishes. pp. 228–235. In: R. Reinboth (ed.) Intersexuality in the Animal Kingdom. Springer-Verlag, Berlin. https://doi.org/10.1007/978-3-642-66069-6_22

Pfennig KS (2007) Facultative mate choice drives adaptive hybridization. Science, 318(5852):965. https://doi.org/10.1126/science.1146035

Phelps SM, Rand AS, Ryan MJ (2006) A cognitive framework for mate choice and species recognition. Am Nat. 167(1):28–42. https://doi.org/10.1086/498538

Plenderleith M, van Oosterhout C, Robinson RL, Turner GF (2005) Female preference for conspecific males based on olfactory cues in a Lake Malawi cichlid fish. Biol Lett. 1(4):411–414. https://doi.org/10.1098/rsbl.2005.0355

Poll M (1986) Classification des cichlidae du Lac Tanganyika: Tribus, genres et espèces. Bull Acad R Belg. 8:1–163.

R core team (2017) R: A Language and environment for statistical computing. R foundation for statistical computing, Vienna, Austria.

Ronco F, Büscher HH, Indermauer A, Salzburger W (2020). The taxonomic diversity of the cichlid fish fauna of ancient Lake Tanganyika, East Africa. J. Great Lakes Res. 46:1067–1078. https://doi.org/10.1016/j.jglr.2019.05.009

Schneider CA, Rasband WS, Eliceiri KW (2012) NIH Image to ImageJ: 25 years of image analysis. Nat Met. 9(7):671–675. https://doi.org/10.1038/nmeth.2089

Roux C, Fraïsse C, Romiguier J, Anciaux Y, Galtier N, Bierne N (2016) Shedding light on the grey zone of speciation along a continuum of genomic divergence. PLOS Biol. 14(12):e2000234. https://doi.org/10.1371/journal.pbio.2000234

Rüber L, Meyer A, Sturmbauer C, Verheyen E (2001). Population structure in two sympatric species of the Lake Tanganyika cichlid tribe Eretmodini: Evidence for introgression. Mol Ecol. 10(5):1207–1225. https://doi.org/10.1046/j.1365-294x.2001.01259.x

Salzburger W (2018). Understanding explosive diversification through cichlid fish genomics. Nat Rev Genet. 19:705–717. https://doi.org/10.1038/s41576-018-0043-9

Salzburger W, Braasch I, Meyer A (2007). Adaptive sequence divergence in a colour gene involved in the formation of the characteristic egg-dummies of male Haplochromis cichlid fishes. BMC Biol. 5(51):13pp. https://doi.org/10.1186/1741-7007-5-51

Seehausen O (2004) Hybridisation and adaptive radiation. Trends Ecol Evol. 19(4):198–207. https://doi.org/10.1016/j.tree.2004.01.003

Seehausen O, van Alphen JJM (1998) The effect of male coloration on female mate choice in closely related Lake Victoria cichlids *(Haplochromis nyererei* complex). Behav Ecol Sociobiol. 42:1–8. https://doi.org/

Seehausen O, Terai Y, Magalhaes IS, Carleton KL, Mrosso HDJ, Miyagi R, van der Sluijs I, Schneider MV, Maan ME, Tachida H, Imai H, Okada N (2008) Speciation through sensory drive in cichlid fish. Nature. 455:620–626. https://doi.org/10.1038/nature07285

Sefc KM, Mattersdorfer K, Ziegelbecker A, Neuhüttler N, Steiner O, Goessler W, Koblmüller S (2017) Shifting barriers and phenotypic diversification by hybridisation. Ecol Lett. 20(5):651–662. https://doi.org/10.1111/ele.12766

Sommer-Trembo C, Plath M, Gismann J, Helfrich C, Bierbach D (2017) Context-dependent female mate choice maintains variation in male sexual activity. R Soc Open Sci. 4:170303. https://doi.org/10.1098/rsos.170303

Strecker U, Kodric-Brown A (1999) Mate recognition systems in a species flock of Mexican pupfish. J Evolution Biol. 12(5):927–935. https://doi.org/10.1046/j.1420-9101.1999.00096.x

Sturmbauer C, Fuchs C, Harb G, Damm E, Duftner N, Maderbacher M, Koch M, Koblmüller S (2008) Abundance, distribution, and territory areas of rock-dwelling Lake Tanganyika cichlid fish species. Hydrobiologia. 615(1):57–68. https://doi.org/10.1007/978-1-4020-9582-5_5

Sullivan BK (2009) Mate recognition, species boundaries and the fallacy of “species recognition”. Open Zool. 2:86–90. https://doi.org/10.2174/1874336601002009086

Svensson EI, Karlsson K, Friberg M, Eroukhmanoff F (2007) Gender differences in species recognition and the evolution of asymmetric sexual isolation. Curr Biol. 17(22):1943–1947. https://doi.org/10.1016/j.cub.2007.09.038

Tinbergen N (1948) Social releasers and the experimental method required for their study. Wilson Bull. 60(1):6–51.

Turner GF, Seehausen O, Knight ME, Allender CJ, Robinson RL (2001) How many species of cichlid fishes are there in African lakes. Mol Ecol. 10(3):793–806. https://doi.org/10.1046/j.1365-294x.2001.01200.x

Theis A, Salzburger W, Egger B (2012) The function of anal fin egg-spots in the cichlid fish Astatotilapia burtoni. PLoS ONE 7(1):e29878. https://doi.org/10.1371/journal.pone.0029878

Thomas GH, Székely T (2005) Evolutionary pathways in shorebird breeding systems: sexual conflict, parental care, and chick development. Evolution 59(10):2222–2230. https://doi.org/10.1111/j.0014-3820.2005.tb00930.x

Trivers RL (1972) Parental investment and sexual selection. in Sexual selection and the descent of man, EBG. Campbell, ed. (136–179). Aldine, Chicago.

Van Oppen MJH, Turner GF, Rico C, Robinson RL, Deutsch JC, Genner MJ, Hewitt GM (1998) Assortative mating among rock-dwelling cichlid fishes supports high estimates of species richness from Lake Malawi. Mol Ecol. 7(8):991–1001. https://doi.org/10.1046/j.1365-294x.1998.00417.x

Verrell PA (1990) Frequency of interspecific mating in salamanders of the plethodontid genus *Desmognathus*: different experimental designs may yield different results. J Zool. 221(3):441–451. https://doi.org/10.1111/j.1469-7998.1990.tb04012.x

Verzijden MN, Korthof REM, ten Cate C (2008) Females learn from mothers and males learn from others. The effect of mother and siblings on the development of female mate preferences and male aggression biases in Lake Victoria cichlids, genus *Mbipia*. Behav Ecol Sociobiol. 62: 1359–1368. https://doi.org/10.1007/s00265-008-0564-x

Verzijden MN, ten Cate, C (2007) Early learning influences species assortative mating preferences in Lake Victoria cichlid fish. Biol lett. 3(2): 134–136. https://doi.org/10.1098/rsbl.2006.0601

Werner NY, Lotem A (2003) Choosy males in a haplochromine cichlid: first experimental evidence for male mate choice in a lekking species. Anim Behav. 66(2):293–298. https://doi.org/10.1006/anbe.2003.2208

Wirtz, P (1999) Mother species-father species: unidirectional hybridization in animals with female choice. Animal Behav. 58:1–12. https://doi.org/10.1006/anbe.1999.1144.

Wong BMB, Jennions MD (2003) Costs influence male mate choice in a freshwater fish. P Roy Soc B-Biol Sci. 270(Suppl 1):S36–S38. https://doi.org/10.1098/rsbl.2003.0003

Zoppoth P, Koblmüller S, Sefc KM (2013) Male courtship preferences demonstrate discrimination against allopatric colour morphs in a cichlid fish. J Evolution Biol. 26(3):577–586. https://doi.org/10.1111/jeb.12074

