## Supplement 1 for "The initial response of females towards congeneric males matches the propensity to hybridise in *Ophthalmotilapia*"

| Specimen | Date | Tank | Focal specimen | Side Focal | Weight focal | Gonadal stage | Non-focal specimen | Weight non-focal | Comment |
|---|---|---|---|---|---|---|---|---|---|
| ON21 | 09 Feb 2016 | 1 | O. nasuta floater male | Left | 12.7 | testes | O. nasuta male | 20.7 | Not used |
| ON22 | 09 Feb 2016 | 2 | O. nasuta female | Right | 16 | 4 | O. nasuta female | | |
| ON23 | 09 Feb 2016 | 3 | O. nasuta female | Right | 19.8 | 3 | O. ventralis male | 10.28 | |
| ON24 | 10 Feb 2016 | 1 | O. nasuta female | Left | 19.5 | 5 | No fish | / | |
| ON26 | 11 Feb 2016 | 1 | O. nasuta floater male | left | 19.4 | testes | O. ventralis male | 15.3 | Not used |
| ON27 | 11 Feb 2016 | 2 | O. nasuta female | Right | 15.4 | 5 | O. nasuta male | 12.7 | |
| ON28 | 11 Feb 2016 | 3 | O. nasuta female | Right | 11 | 3 | O. nasuta female | 6.16 | |
| ON30 | 12 Feb 2016 | 3 | O. nasuta female | Right | 12.92 | 5 | No fish | / | |
| ON31 | 13 Feb 2016 | 1 | O. nasuta female | Left | 11 | 4 | O. nasuta female | | |
| ON32 | 13 Feb 2016 | 2 | O. nasuta floater male | Right | 10.5 | testes | O. ventralis male | 10.01 | Not used |
| ON33 | 13 Feb 2016 | 3 | O. nasuta floater male | Right | 9.55 | testes | O. nasuta male | 20.1 | Not used |
| ON34 | 14 Feb 2016 | 1 | O. nasuta female | left | 9.88 | 3 | O. ventralis male | 15.4 | |
| ON35 | 14 Feb 2016 | 2 | O. nasuta floater male | Right | 7.8 | | No fish | / | Not used |
| ON36 | 14 Feb 2016 | 3 | O. nasuta female | Right | 6.7 | 5 | O. nasuta male | 20.7 | |
| ON37 | 15 Feb 2016 | 1 | O. nasuta female | Left | 8.7 | 5 | O. nasuta male | 20.7 | |
| ON38 | 15 Feb 2016 | 2 | O. nasuta female | Right | 6.76 | 3 | O. ventralis male | 10.1 | Not used |
| ON39 | 15 Feb 2016 | 3 | O. nasuta female | Right | 6.16 | 3 | No fish | / | |
| ON41 | 16 Feb 2016 | 1 | O. nasuta female | left | 11.8 | 4 | O. nasuta male | 12.7 | |
| ON43 | 16 Feb 2016 | 3 | O. nasuta floater male | Right | 12.6 | testes | O. ventralis male | 10.8 | Not used |
| ON44 | 28 Aug 2018 | 1 | O. nasuta female | Right | 19 | 4 | O. ventralis male | 19 | |
| ON45 | 28 Aug 2018 | 2 | O. nasuta female | Left | 11 | 4 | No fish | / | |
| ON46 | 28 Aug 2018 | 3 | O. nasuta female | Left | 30 | 3 | O. nasuta male | 19 | |
| ON47 | 29 Aug 2018 | 1 | O. nasuta female | Right | 11 | 5 | No fish | / | |
| ON48 | 29 Aug 2018 | 2 | O. nasuta female | Left | 17 | 5 | O. ventralis male | 12 | |
| ON49 | 29 Aug 2018 | 3 | O. nasuta female | Left | 13 | 5 | O. nasuta female | 11 | |
| ON50 | 30 Aug 2018 | 1 | O. nasuta female | Right | 7 | 4 | O. nasuta female | 15 | |
| ON51 | 30 Aug 2018 | 2 | O. nasuta female | Left | 17 | 4 | O. ventralis male | 19 | Not used |
| OV13 | 10 Oct 2018 | 1 | O. ventralis female | Left | 14 | 2 | O. ventralis female | 10 | |
| OV14 | 10 Oct 2018 | 2 | O. ventralis female | Right | 16 | 2 | O. ventralis male | 16 | |
| OV15 | 10 Oct 2018 | 3 | O. ventralis female | Left | 12 | 5 | O. nasuta male | 17 | |
| OV16 | 11 Oct 2018 | 1 | O. ventralis female | Right | 10 | 2 | O. nasuta male | 16 | |
| OV17 | 11 Oct 2018 | 2 | O. ventralis female | Left | 12 | 3 | No fish | | |
| OV18 | 11 Oct 2018 | 3 | O. ventralis female | Right | 10 | 4 | O. ventralis male | 17 | |
| OV19 | 12 Oct 2018 | 1 | O. ventralis female | Left | 11 | 1 | O. ventralis male | 16 | |
| OV20 | 12 Oct 2018 | 2 | O. ventralis female | Right | 8 | 3 | O. ventralis female | 10 | |
| OV21 | 12 Oct 2018 | 3 | O. ventralis female | Right | 13 | 4 | No fish | / | |
| OV22 | 15 Oct 2018 | 1 | O. ventralis female | Left | 9 | 4 | No fish | / | |
| OV23 | 15 Oct 2018 | 2 | O. ventralis female | Left | 10 | 4 | O. nasuta male | 42 | |
| OV24 | 15 Oct 2018 | 3 | O. ventralis female | Right | 8 | 5 | O. ventralis male | 17 | |
| OV25 | 16 Oct 2018 | 1 | O. ventralis female | Right | 9 | 5 | O. ventralis male | 17 | |
| OV26 | 16 Oct 2018 | 2 | O. ventralis female | Right | 10 | 3 | No fish | / | |
| OV27 | 16 Oct 2018 | 3 | O. ventralis female | Left | 14 | 5 | O. ventralis female | 10 | |
| OV28 | 17 Oct 2018 | 1 | O. ventralis female | Right | 12 | 4 | O. ventralis female | 9 | |
| OV29 | 17 Oct 2018 | 2 | O. ventralis female | Right | 9 | 5 | O. nasuta male | 18 | |
| OV30 | 17 Oct 2018 | 3 | O. ventralis female | Left | 9 | 5 | No fish | / | |
| OV31 | 18 Oct 2018 | 1 | O. ventralis female | Left | 9 | 4 | O. nasuta male | 22 | |
| OV32 | 18 Oct 2018 | 2 | O. ventralis female | Left | 6 | 3 | O. ventralis female | 8 | |
| OV33 | 18 Oct 2018 | 3 | O. ventralis female | Right | 10 | 4 | O. nasuta male | 38 | |
