## Supplementary figures and images for "The initial response of females towards congeneric males matches the propensity to hybridise in *Ophthalmotilapia*"

### Supplement 2

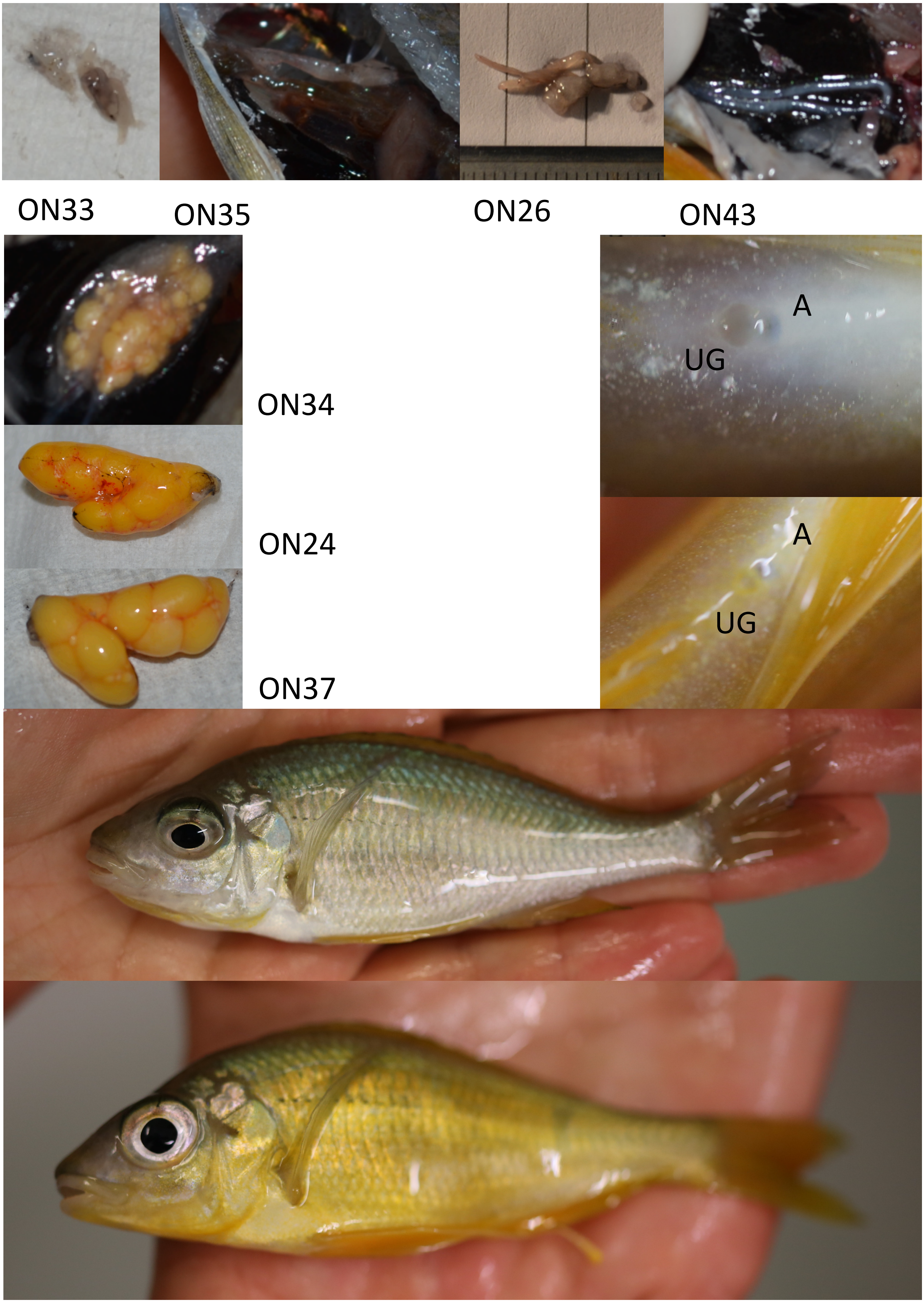

### Supplement 3

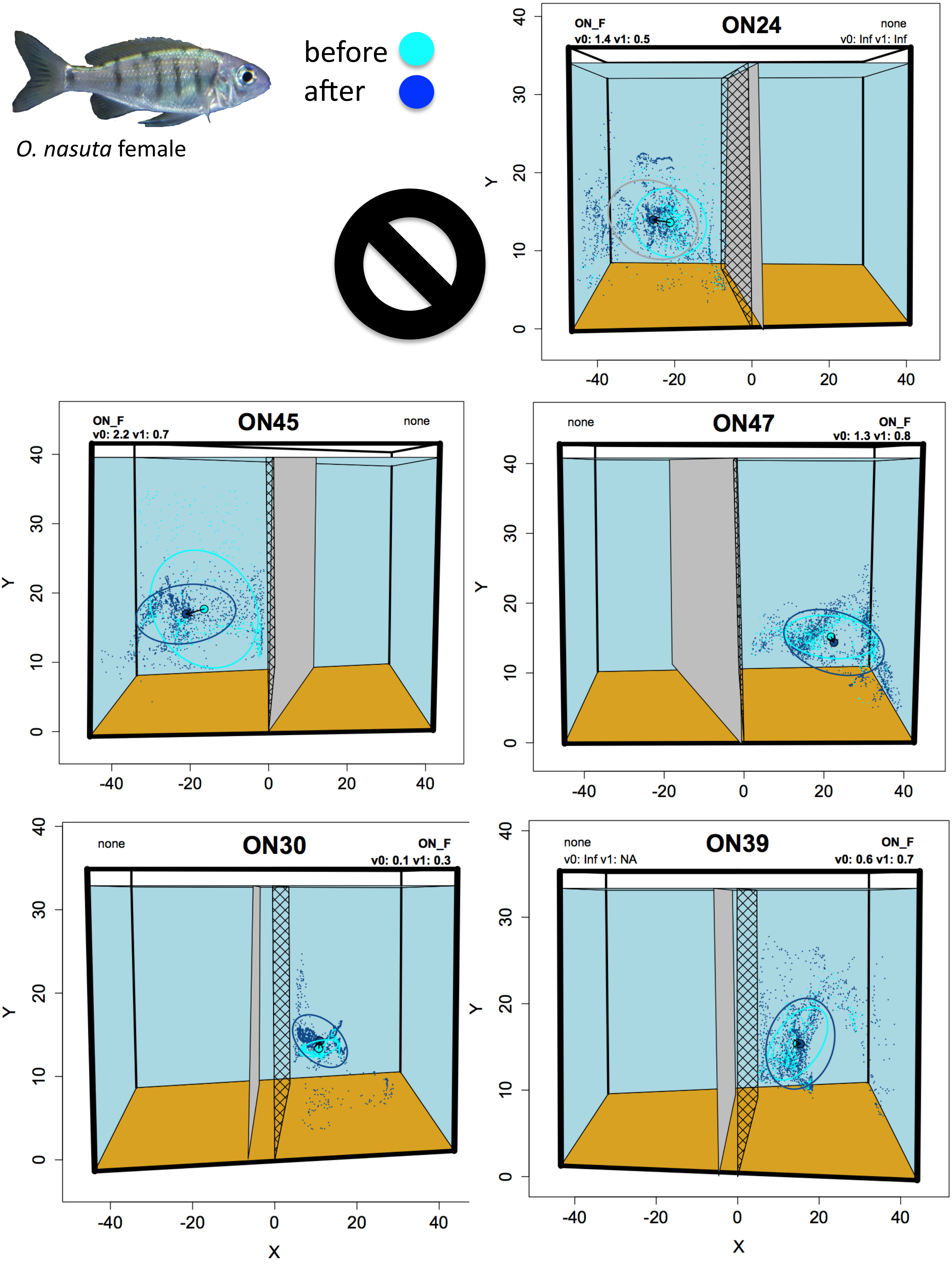

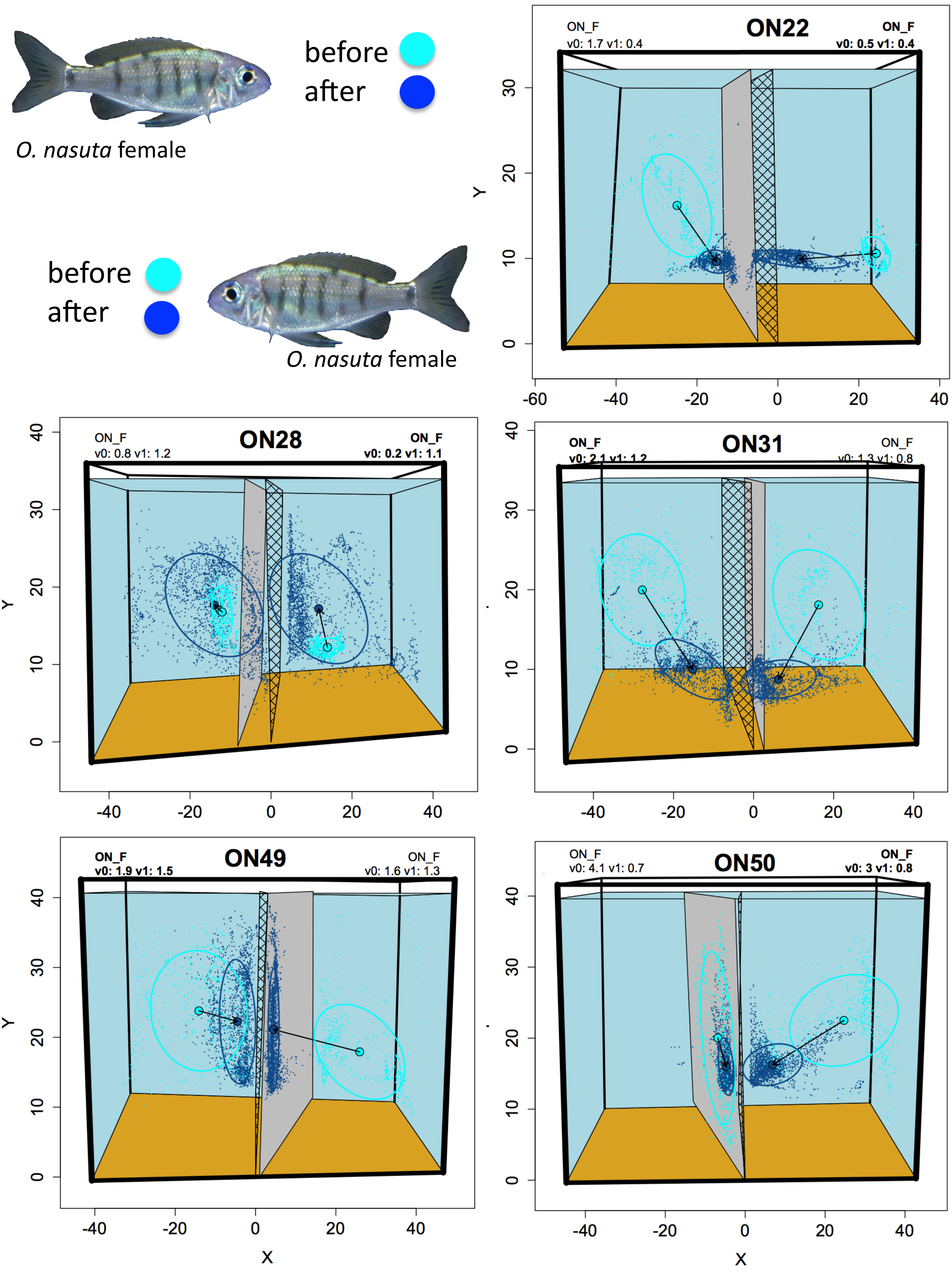

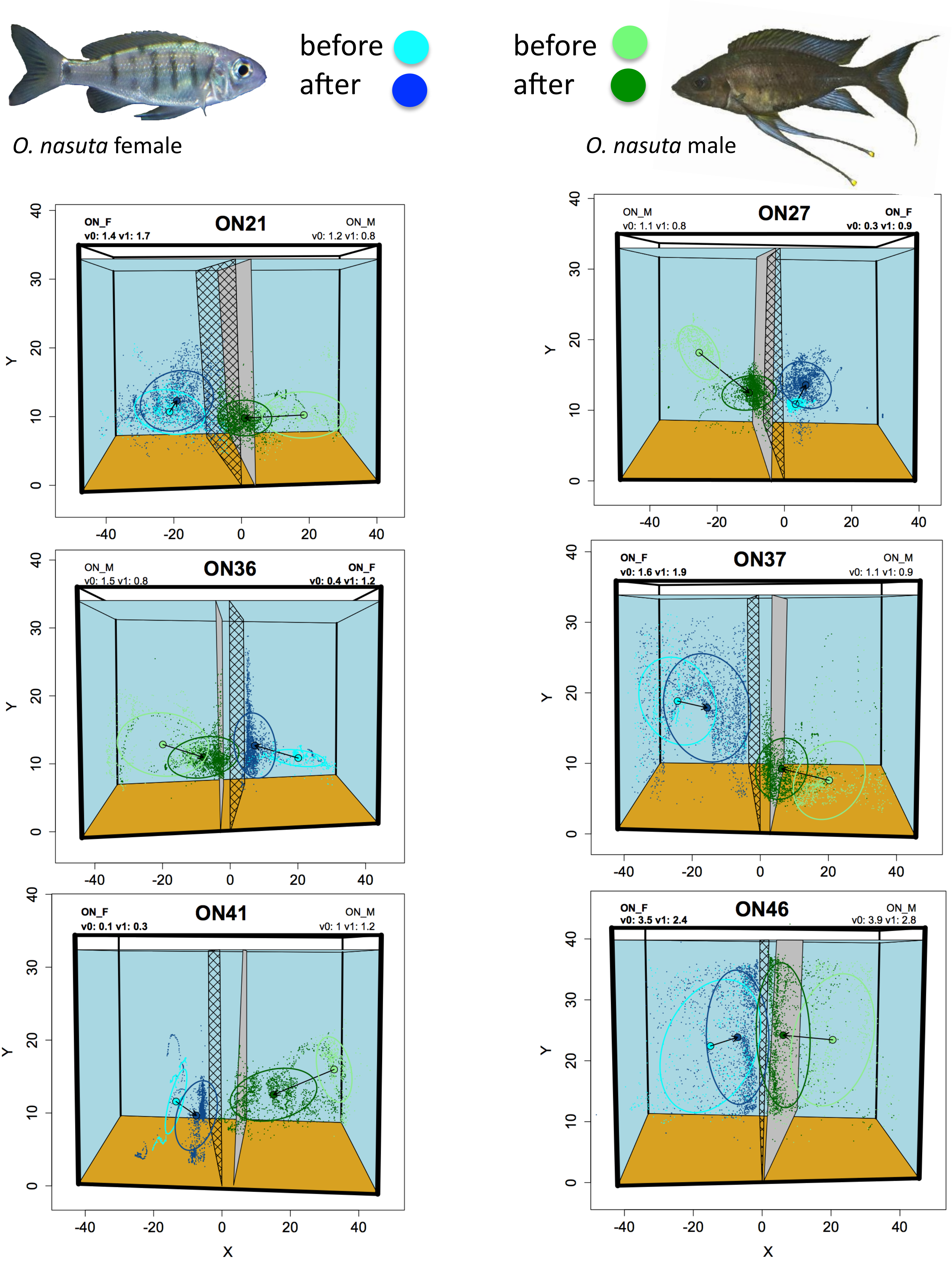

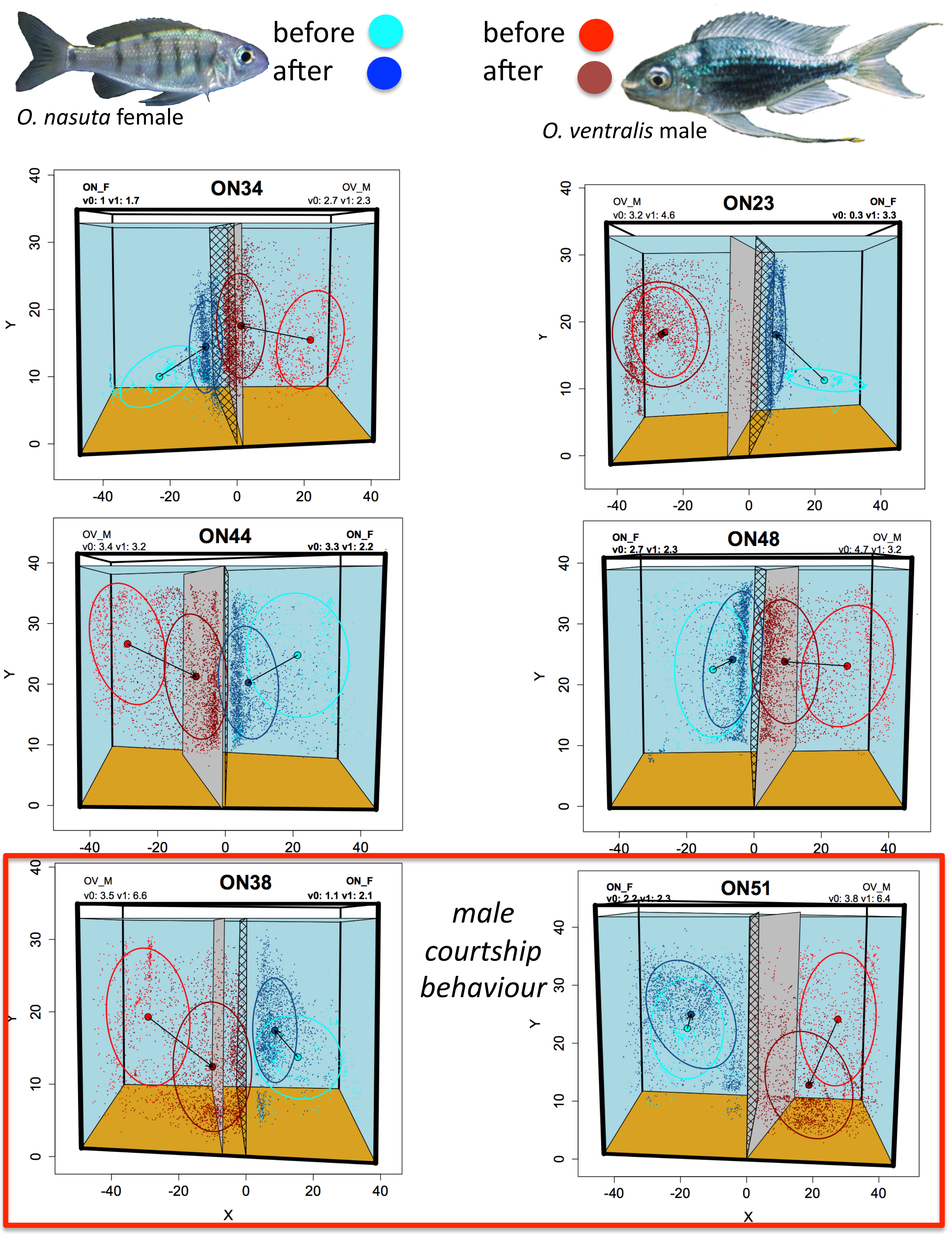

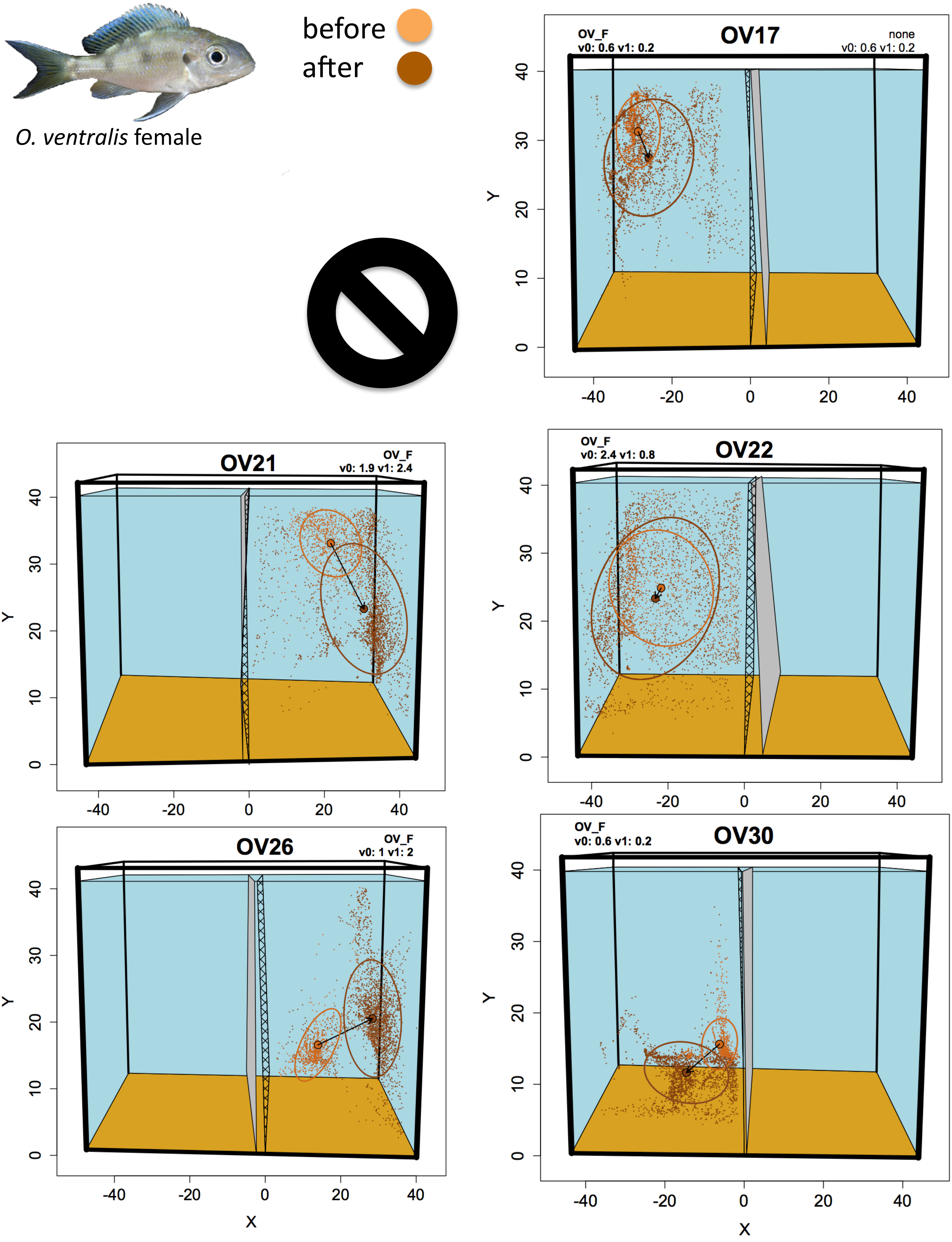

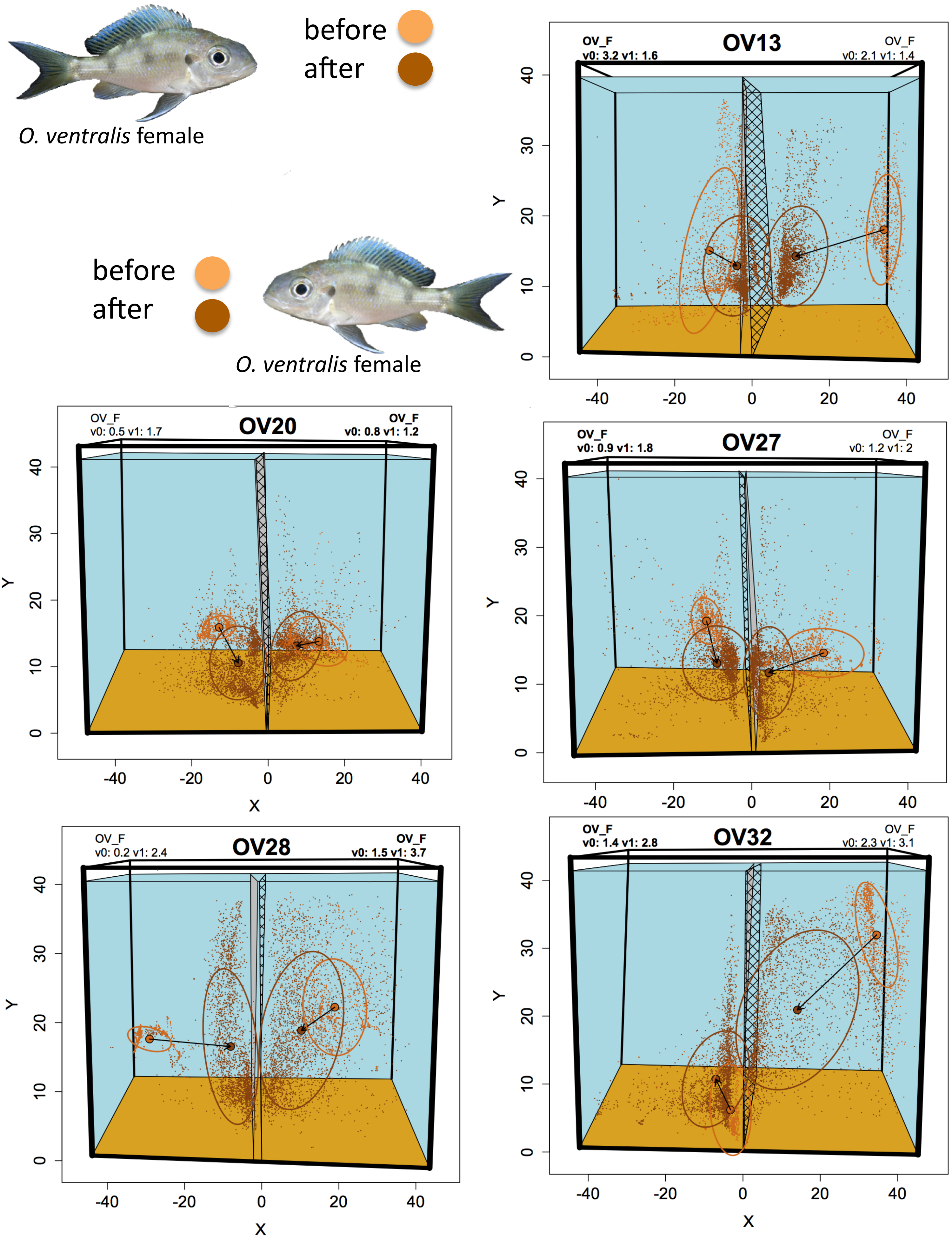

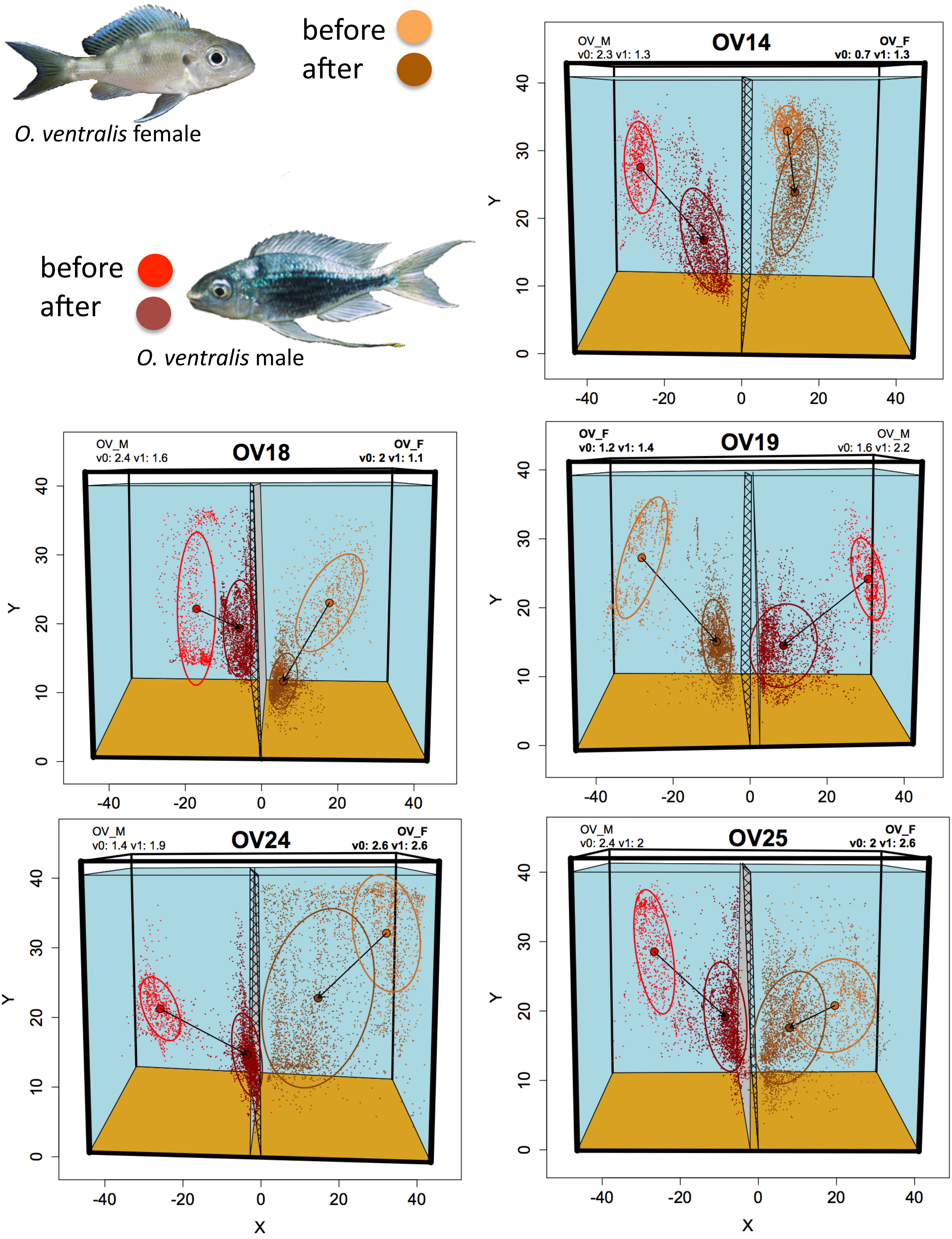

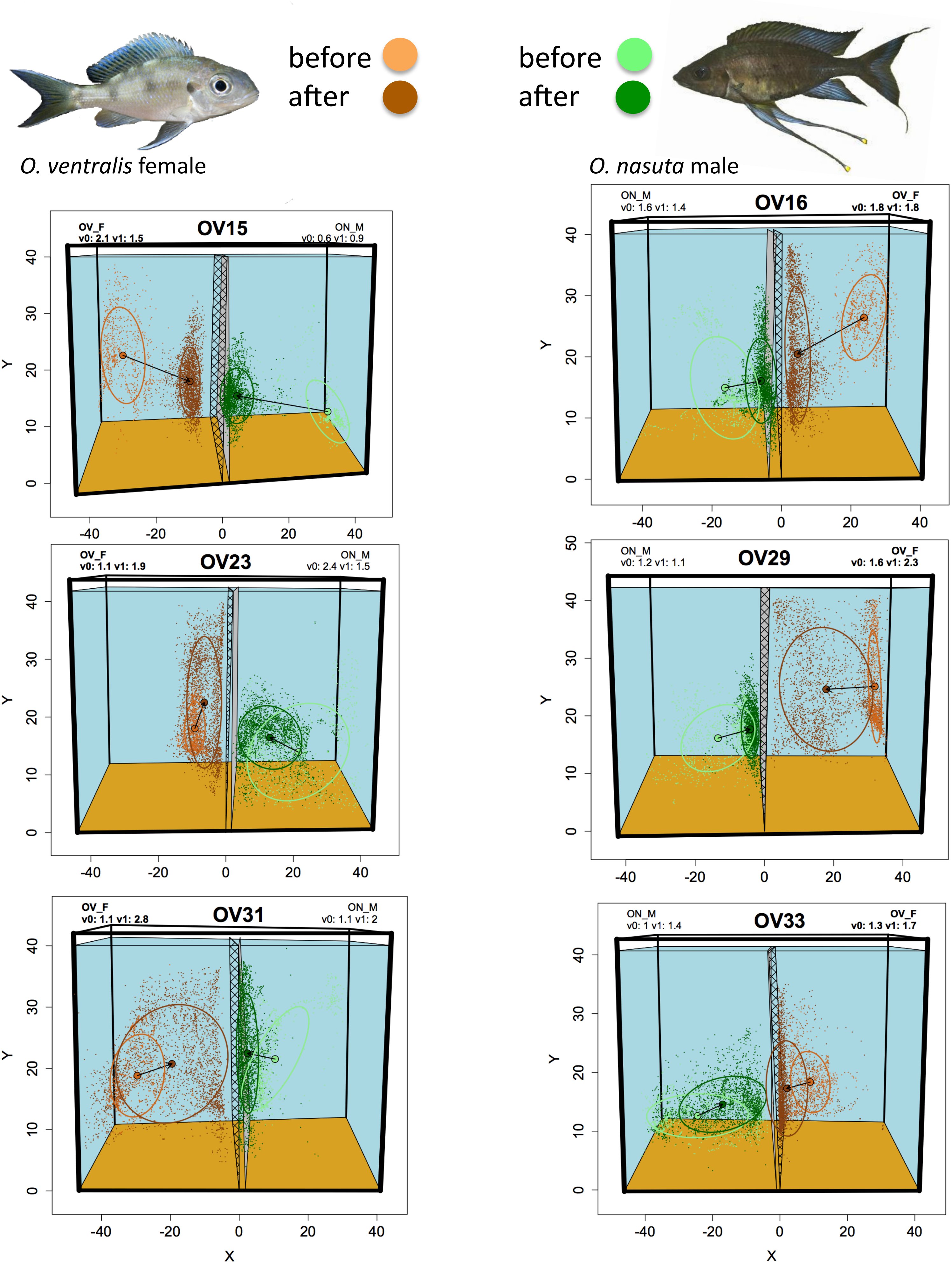

### Supplement 6

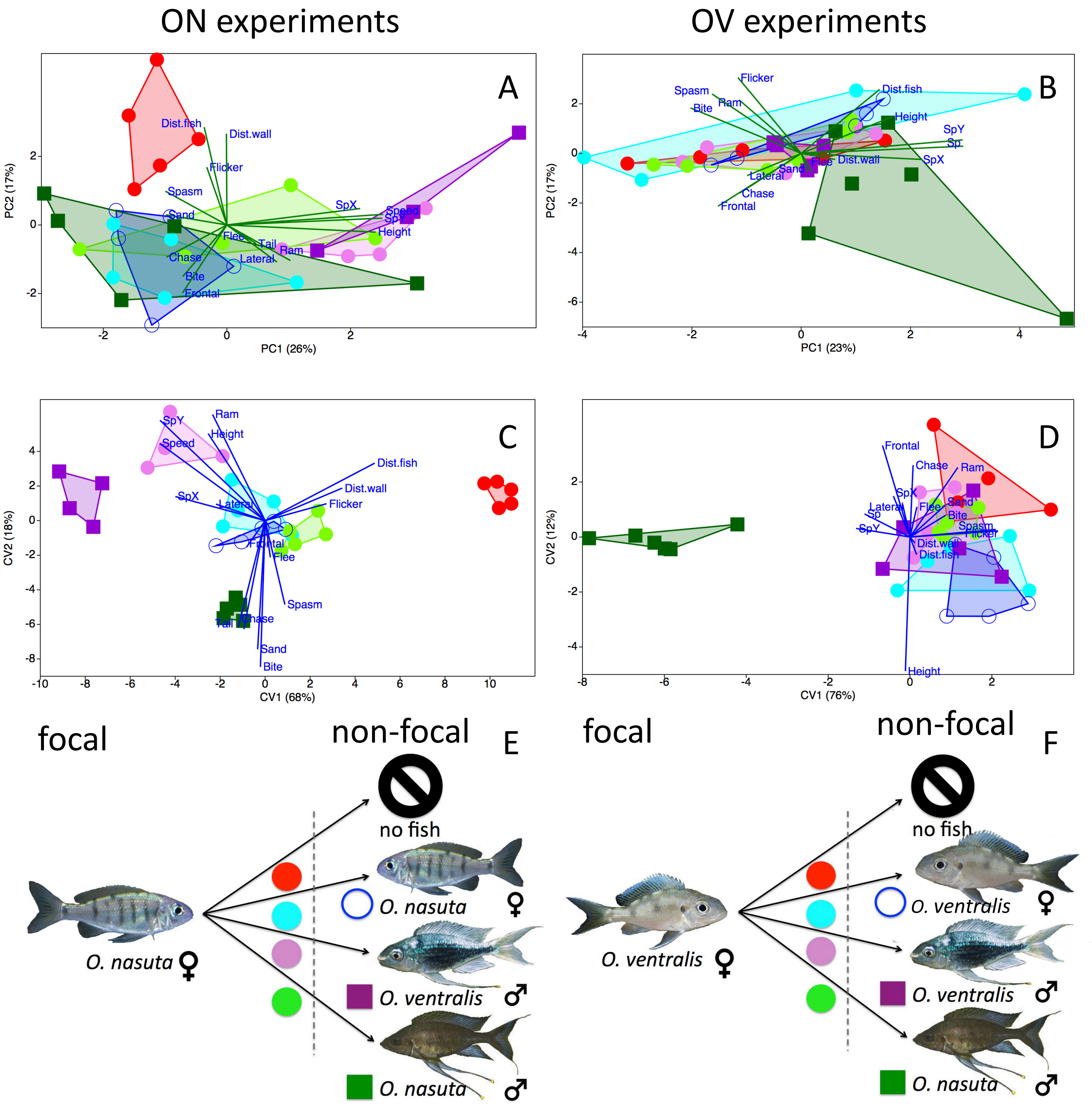

### Supplement 7

ON experiments Focal ON experiments Non-Focal

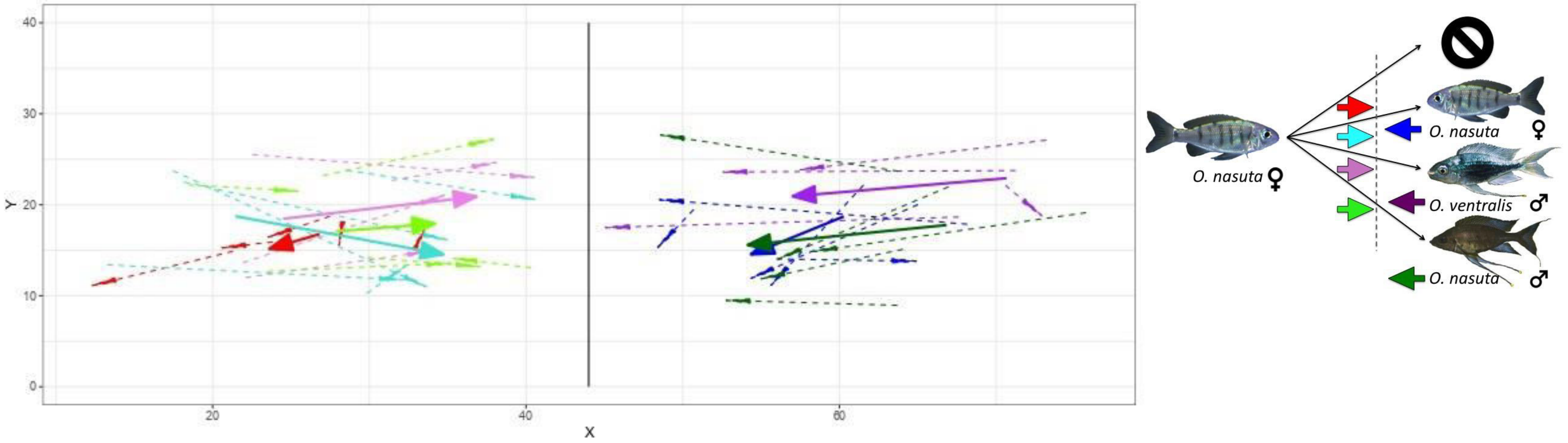


OV experiments Focal OV experiments Non-Focal

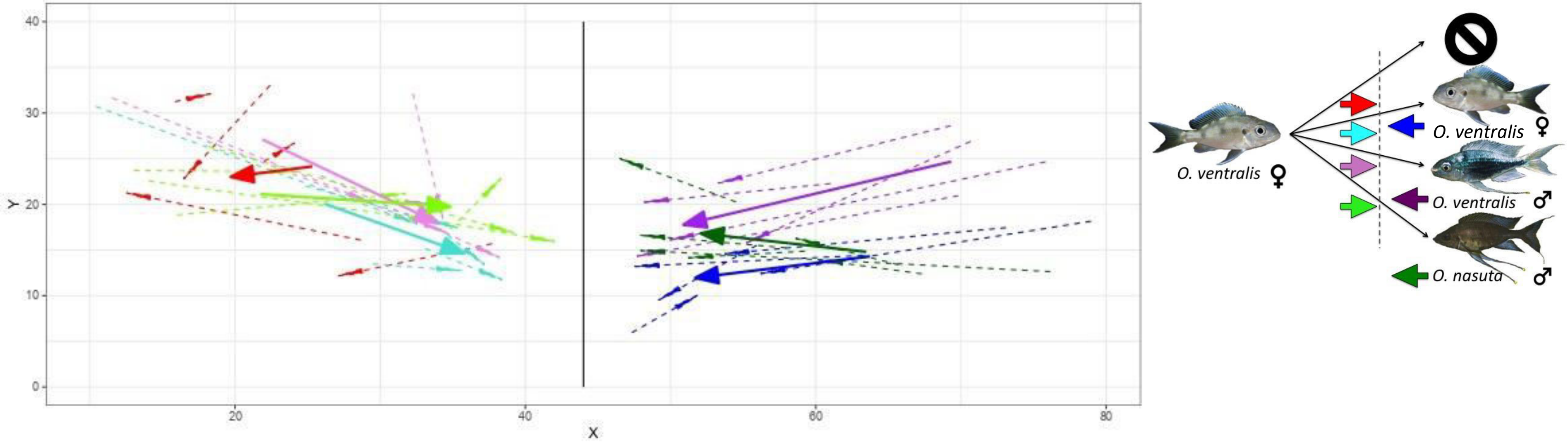
