## Supplement 5 for "The initial response of females towards congeneric males matches the propensity to hybridise in *Ophthalmotilapia*"

**Supplement 5.1.** Loadings and explained variance of the PCAs performed on the behavioural parameters collected during three time intervals (Before, After1, After2) for the ON and OV experiments.

| | ON: Before | | ON: After1 | | ON: After2 | | OV: Before | | OV: After1 | | OV: After2 | |
|---|---|---|---|---|---|---|---|---|---|---|---|---|
| | **PC1** | **PC2** | **PC1** | **PC2** | **PC1** | **PC2** | **PC1** | **PC2** | **PC1** | **PC2** | **PC1** | **PC2** |
| **ev** | 4.428 | 2.018 | 3.859 | 2.727 | 4.102 | 2.689 | 3.036 | 2.135 | 3.656 | 2.193 | 3.469 | 2.503 |
| **var** | 40.253 | 18.347 | 24.116 | 17.041 | 25.635 | 16.803 | 25.297 | 17.793 | 24.372 | 14.623 | 23.124 | 16.687 |
| **Chase** | / | / | -0.173 | 0.302 | -0.180 | -0.174 | / | / | -0.245 | 0.192 | -0.175 | -0.248 |
| **Flee** | 0.350 | 0.217 | 0.146 | 0.114 | 0.063 | -0.080 | 0.172 | -0.004 | 0.048 | 0.173 | 0.033 | -0.119 |
| **Lateral** | / | / | 0.084 | 0.298 | 0.149 | -0.200 | / | / | -0.253 | 0.359 | -0.153 | -0.139 |
| **Frontal** | / | / | -0.077 | 0.416 | -0.133 | -0.367 | / | / | -0.326 | 0.270 | -0.236 | -0.330 |
| **Bite** | / | / | -0.128 | 0.305 | -0.132 | -0.277 | -0.083 | 0.278 | -0.321 | 0.282 | -0.314 | 0.285 |
| **Ram** | 0.273 | 0.125 | 0.184 | 0.163 | 0.190 | -0.192 | 0.032 | 0.494 | -0.116 | 0.321 | -0.176 | 0.335 |
| **Sand** | -0.084 | -0.006 | -0.213 | -0.187 | -0.182 | 0.055 | -0.132 | 0.345 | -0.022 | -0.182 | 0.019 | -0.118 |
| **Spasm** | -0.022 | 0.479 | -0.124 | -0.148 | -0.184 | 0.182 | -0.063 | -0.252 | -0.138 | -0.018 | -0.253 | 0.372 |
| **Tail** | / | / | 0.036 | 0.093 | 0.087 | -0.100 | / | / | / | / | / | / |
| **Flicker** | -0.161 | 0.518 | -0.170 | -0.117 | -0.060 | 0.311 | -0.083 | -0.023 | -0.013 | -0.212 | -0.180 | 0.475 |
| **Dist.wall** | 0.179 | -0.429 | -0.040 | -0.436 | -0.002 | 0.491 | 0.294 | -0.223 | 0.233 | -0.268 | 0.096 | -0.038 |
| **Dist.fish** | 0.128 | -0.468 | -0.101 | -0.459 | -0.068 | 0.526 | 0.253 | -0.347 | 0.213 | -0.088 | 0.223 | 0.401 |
| **Sp** | 0.451 | 0.112 | 0.470 | -0.100 | 0.466 | 0.057 | 0.528 | 0.204 | 0.413 | 0.388 | 0.460 | 0.045 |
| **SpX** | 0.439 | 0.048 | 0.391 | -0.082 | 0.397 | 0.089 | 0.372 | 0.405 | 0.318 | 0.324 | 0.339 | -0.037 |
| **SpY** | 0.413 | 0.141 | 0.468 | -0.104 | 0.464 | 0.037 | 0.524 | 0.064 | 0.405 | 0.370 | 0.458 | 0.085 |
| **Height** | 0.393 | -0.024 | 0.436 | 0.080 | 0.444 | -0.038 | 0.304 | -0.340 | 0.310 | 0.013 | 0.258 | 0.229 |

**Supplement 5.2.** Loadings and explained variance of the CVAs performed on the behavioural parameters collected during three time intervals (Before, After1, After2) for the ON and OV experiments.

| | ON: Before | | ON: After1 | | ON: After2 | | OV: Before | | OV: After1 | | OV: After2 | |
|---|---|---|---|---|---|---|---|---|---|---|---|---|
| | **CV1** | **CV2** | **CV1** | **CV2** | **CV1** | **CV2** | **CV1** | **CV2** | **CV1** | **CV2** | **CV1** | **CV2** |
| **ev** | 6.245 | 1.522 | 18.534 | 6.544 | 34.783 | 9.049 | 2.952 | 2.317 | 7.480 | 2.751 | 9.172 | 1.435 |
| **var** | 66.710 | 16.260 | 57.420 | 20.270 | 67.910 | 17.670 | 47.970 | 37.660 | 61.260 | 22.530 | 75.670 | 11.840 |
| **Chase** | / | / | -0.037 | -0.093 | -0.023 | -0.152 | / | / | -0.096 | -0.088 | 0.028 | -0.957 |
| **Flee** | 0.191 | -0.048 | -0.047 | -0.058 | 0.006 | -0.052 | -0.079 | -0.011 | -0.017 | -0.208 | 0.038 | -0.249 |
| **Lateral** | / | / | -0.034 | -0.024 | -0.053 | 0.023 | / | / | -0.027 | -0.052 | -0.013 | -0.084 |
| **Frontal** | / | / | -0.007 | 0.040 | -0.020 | -0.037 | / | / | 0.001 | -0.131 | -0.118 | -0.585 |
| **Bite** | / | / | -0.017 | -0.232 | -0.005 | -0.205 | 0.154 | -0.022 | -0.079 | -0.037 | 0.226 | -0.205 |
| **Ram** | -0.093 | 0.081 | -0.021 | 0.115 | -0.057 | 0.149 | 0.482 | 0.036 | -8.836 | 0.468 | 15.582 | -33.722 |
| **Sand** | -0.026 | 0.359 | -0.026 | -0.235 | -0.008 | -0.181 | 0.307 | -0.045 | -0.032 | 0.057 | 0.199 | -0.267 |
| **Spasm** | 0.067 | 0.326 | -0.011 | -0.158 | 0.021 | -0.118 | 0.152 | 0.077 | -0.195 | 0.342 | 0.981 | -0.128 |
| **Tail** | / | / | -0.094 | 0.054 | -0.029 | -0.149 | / | / | / | / | / | / |
| **Flicker** | -0.035 | -0.068 | -0.019 | -0.166 | 0.066 | 0.024 | 0.038 | -0.006 | -0.016 | 0.029 | 0.179 | -0.019 |
| **Dist.wall** | 0.179 | 0.243 | 0.073 | 0.014 | 0.083 | 0.046 | -1.152 | 1.353 | 4.367 | 4.229 | 0.558 | 1.016 |
| **Dist.fish** | 0.062 | -0.056 | 0.128 | 0.016 | 0.118 | 0.081 | -0.036 | 1.884 | -0.110 | 0.364 | 0.074 | 0.292 |
| **Sp** | 0.200 | 0.052 | -0.169 | 0.098 | -0.114 | 0.108 | -0.073 | 0.124 | 0.045 | -0.096 | -0.084 | -0.064 |
| **SpX** | 0.162 | 0.079 | -0.174 | 0.064 | -0.097 | 0.034 | 0.061 | 0.044 | 0.008 | -0.037 | -0.016 | -0.058 |
| **SpY** | 0.218 | 0.032 | -0.149 | 0.109 | -0.113 | 0.141 | -0.140 | 0.116 | 0.047 | -0.082 | -0.086 | -0.020 |
| **Height** | 0.149 | -0.005 | -0.082 | 0.041 | -0.062 | 0.122 | -3.494 | 7.228 | 1.603 | 3.055 | -0.133 | 5.773 |
